# Modular Integration of Auditory Instructions and Visual Cues into the Cortical Reach Network

**DOI:** 10.64898/2026.01.20.700205

**Authors:** Gaelle N. Luabeya, Ada Le, Amir Hossein Ghaderi, Lina Musa, Simona Monaco, Erez Freud, J. Douglas Crawford

**Affiliations:** Centre for Integrative and Applied Neuroscience, York University, Canada; Centre for Vision Research, and Vision: Science to Applications Program, York University, Canada; Connected Minds Program, York University, Canada; Department of Biology, York University, Canada; Department of Psychology, York University, Canada; Department of Kinesiology & Health Sciences, York University, Canada; Center for Affective Neuroscience, Development, Learning and Education (CANDLE), University of Southern California, Los Angeles, CA, USA; Center for Mind/Brain Sciences, University of Trento, Italy

**Keywords:** Functional Connectivity, Functional Magnetic Resonance Imaging, Graph Theory Analysis, Multisensory Integration, Reach-to-Grasp

## Abstract

Humans often need to integrate ‘bottom-up’ stimulus attributes (e.g., object shape) with ‘top-down’ instructions (e.g., how to pick it up) for adaptive behavior. To investigate how such cues are integrated into the cortical reach network, we employed an fMRI paradigm where participants viewed a cube to their left or right (visual cue) and were verbally instructed to use a horizontal or vertical grip (auditory cue), in varying order. Univariate voxel-wise and region-of-interest analysis confirmed the expected order-dependent activation of sensory regions, followed by robust parietofrontal activation. Graph-theory analysis revealed two significant subnetworks (modules): an Occipital-Parietal module (likely integrating the visual cue into the reach plan) and a Temporal-Frontal module (likely integrating the auditory cue into the grasp plan), both converging in somatomotor cortex. Cue-order influenced modularity, i.e., the module corresponding to the first cue tended to dominate contralateral motor cortex, and conversely a classification model decoded cue-order based on modularity. Local network hubs were order-independent and occupied traditional reach areas, whereas global (between module) hubs were order-dependent and more distributed. Overall, these results suggest that multisensory integration of top-down and bottom-up cues for action is a network phenomenon that relies on both lateral and serial communication between parallel sensorimotor modules.

## 1. INTRODUCTION

Voluntary actions are driven by both ‘bottom-up’ sensory cues and ‘top-down’ cognitive cues for motor planning (Ariani et al. 2015; Fried et al. 2017; Virameteekul and Bhidayasiri 2022). For example, the first time one uses a fork, the visual system provides information about its shape, size, orientation, and location, but a caregiver likely provides instructions on how to grasp it correctly. In such cases, the brain must thus integrate both bottom-up visual cues and top-down instructions to form the optimal motor plan. Although numerous studies have considered that cortical regions support various aspects of action planning and execution (e.g., Glover, 2004; Majdandžić et al., 2007; Gallivan et al., 2011), most such studies did not consider the multisensory-motor integration processes required to integrate their top-down task instructions with their bottom-up sensory stimuli.

Prehension can be broken down into the transport component, which moves the hand to the correct location, and the grasp component, which shapes the hand to the object (Jeannerod 1984; Desmurget et al. 1998). The bottom-up source of visual information for such movements, e.g., location, shape, size, and orientation of the fork described above, can be reliably attributed to well-studied processes in the visual system and involved in the ‘dorsal stream’ parietofrontal circuits for reach (i.e., Goodale and Milner, 1992; Henriques et al., 1998; Crawford et al., 2011; Chen et al., 2023; Luabeya et al., 2024). In monkeys and humans, location information appears to be processed through the ‘dorsal visual stream’, including connections spanning early visual cortex (V1), dorsal occipital cortex, superior parietal-occipital cortex (SPOC), posterior intraparietal cortex (pIPS), and dorsal premotor cortex (PMd) (Mishkin and Ungerleider 1982; Rizzolatti and Matelli 2003; Vesia and Crawford 2012; Gallivan and Culham 2015; Baltaretu et al. 2020). In contrast, grasp involves both ventral and dorsal stream processes, including lateral occipital cortex (LOC), anterior intraparietal cortex (aIPS), and ventral premotor cortex (PMv) (Begliomini et al. 2008; Jacobs et al. 2010; Klein et al. 2023). Ultimately, these high-level planning areas are thought to activate neurons in primary somatosensory (S1) and primary motor (M1) areas, leading to the activation of specific muscle groups for action (Schieber 2001; Umeda et al. 2019; Ariani et al. 2022).

The influence of top-down verbal instructions, although often used in laboratory paradigms, is less understood, perhaps because it involves aspects of cognition traditionally considered outside the realm of sensorimotor control. As noted above, this often involves instructions on how and what to do, especially when learning new tasks. Once learned, this may evolve into an affordance: the implicit knowledge of the various ways an object can be handled (Gibson and Maunsell 1997; Osiurak et al. 2017), but instruction, whether through imitation, signage, or language, continues to play a role in many activities. In the case of verbal instruction, auditory information is thought to be relayed from the primary auditory cortex (A1) through Wernicke’s area to eventually reach frontal regions, facilitating comprehension of the sound (Binder 2000; Hickok and Poeppel 2007; Friederici 2011; Price 2012). A recent resting-state Magnetoencephalography study reported stronger phase coupling between the primary auditory cortex (A1) and the ventral premotor cortex (Bedford et al. 2025), suggesting that sensorimotor networks have a modality-specific organization.

The question remains, how are these two sources of information (top-down verbal instructions for motor strategy and bottom-up sensory features) integrated for planning and execution of reach movements? One possibility is that specific regions of interest (ROIs) in the parietal or frontal cortex perform this function (Miller and Cohen 2001; Andersen and Cui 2009). An alternative approach is to consider this question from a whole-brain network perspective (Musa et al. 2025). Specifically, how are these two types of signals distributed through the cortex during reach-to-grasp planning? A whole-brain network approach also has considerable medical relevance, since stroke and other forms of brain damage can impair cue integration for motor control (Edwards et al. 2019; Guggisberg et al. 2019).

One way to address this question is to phrase it in terms of ‘functional connectivity,’ i.e., correlations between signals across different brain areas (Damoiseaux and Greicius 2009; Honey et al. 2009; Stam et al. 2016). By extracting time series from multiple ROIs as ‘nodes’, and correlating them to construct ‘adjacency matrices’ from all these areas, one can apply network science tools in graph theoretical analysis (GTA) to characterize various properties of their functional network (Bullmore and Sporns 2009; Rubinov and Sporns 2010; Bassett and Sporns 2017). In the field of cognitive neuroscience, GTA has gained popularity in resting-state fMRI (functional Magnetic Resonance Imaging) and EEG (Electroencephalography) studies (Medaglia 2017; Smitha et al. 2017; Sun et al. 2019), but in recent years, it has also been applied to event-related EEG and fMRI data (Ghaderi et al. 2023; Musa et al. 2025; Tomou et al. 2025). With this, one can investigate and compare the global network properties (e.g., energy, efficiency, clustering), intermediate properties (i.e., modularity), and local properties (i.e., hubness) of various functional networks (Newman 2006; Bullmore and Sporns 2009; Rubinov and Sporns 2010; Ghaderi et al. 2018). Task instruction has been shown to alter network properties in the visual system (Musa et al. 2025), but to our knowledge, its influence on multimodal integration in the motor system has not been investigated.

To understand how the brain integrates bottom-up visual cues with top-down verbal instructions for grasp rotation, we employed a cue-separation fMRI task in which participants were visually cued to the object attributes (e.g., location, shape, size, orientation), and verbally instructed on which way to orient the hand (vertical or horizontal grip). In addition, we alternated the order of cue presentation to assess whether the timing of cue presentation influences brain activity and connectivity. From our whole-brain univariate analysis, we expected to see order-dependent activation of the sensory cortex since these responses tend to be transient (Jäncke et al. 2002; Hasson et al. 2008), whereas preparatory motor activity tends to ramp up toward execution (Cappadocia et al. 2017). Alongside voxelwise analyses, we conducted ROI-based analyses to test a priori hypotheses in reach-and-grasp regions, and then a comprehensive GTA network analysis based on the functional connectivity of the BOLD time series.

Based on the literature cited above, we expected that sensory areas (primary visual and auditory cortex) would show order-dependent activation, depending on the order of stimulus cue presentation, but that this order-dependence would successfully decrease as this information was integrated in higher level parietofrontal areas, progressing to motor cortex. In terms of network analysis, one would expect 1) visual activation to correlate with dorsal stream parietal activation, 2) auditory signals in temporal cortex might correlate with activity in higher-level frontal planning areas, and 3) for these two parallel sensorimotor ‘modules’ to converge somehow in somatomotor cortex. One would expect local hubs to emerge within these modules, whereas global ‘between module’ hubs should reveal additional lateral integrative mechanisms. Finally, we used a machine-learning classification technique to assess which network features encode cue order.

## 2. METHODS

### 2.1. Participants

This study was a secondary analysis of a dataset that was only published in abstract form (Le et al. 2015). 12 right-handed individuals (8 females, 4 males; age range, 21-37 years; median, 27 years) participated in the original study, based on the numbers used in contemporaneous sensorimotor imaging studies (Gallivan et al. 2013; Chen et al. 2014; Fabbri et al. 2016; Cappadocia et al. 2017). We acknowledge that these participant numbers are low compared to the current standards for perceptual / cognitive tasks, but reach planning and execution produce highly reliable and robust BOLD activation (Bennett and Miller 2010; Poldrack et al. 2017), such that even modest participant numbers have been shown to provide sufficient power for functional network analysis (Musa et al. 2025). Further, we conducted a post hoc power analysis to confirm that we had sufficient statistical power for analysis of even our least robust region-of-interest results, and used permutation testing and cross-validation to establish the reliability of our GTA findings (see *Statistical Analysis* section, below). Participants exhibited normal or corrected-to-normal visual acuity with no reported neurological disorders and were all right-handed. Participants provided informed consent before the experiment and received monetary compensation for their time. All experimental procedures were approved by the York University Human Participants Review Committee (Certificate #: 2015-219; protocol renewed November 22, 2021).

### 2.2. Apparatus and Stimuli

Data were collected at York University (Toronto, Canada) using a 3-T whole-body MRI system (Siemens Magnetom TIM Trio, Erlangen, Germany). Participants lay in the platform on which the stimuli are presented. Headphones were worn to hear auditory instructions. To reduce motion artifacts, the head was immobilized using foam cushions, and a strap further restricted their upper arms. To facilitate the view of the stimuli in the head-titled setup, the anterior part of the 12-channel coil was removed and replaced by a 4-channel flex coil, while the posterior portion of the 12-channel coil was maintained (Fig. 1A).

**Figure 1.**
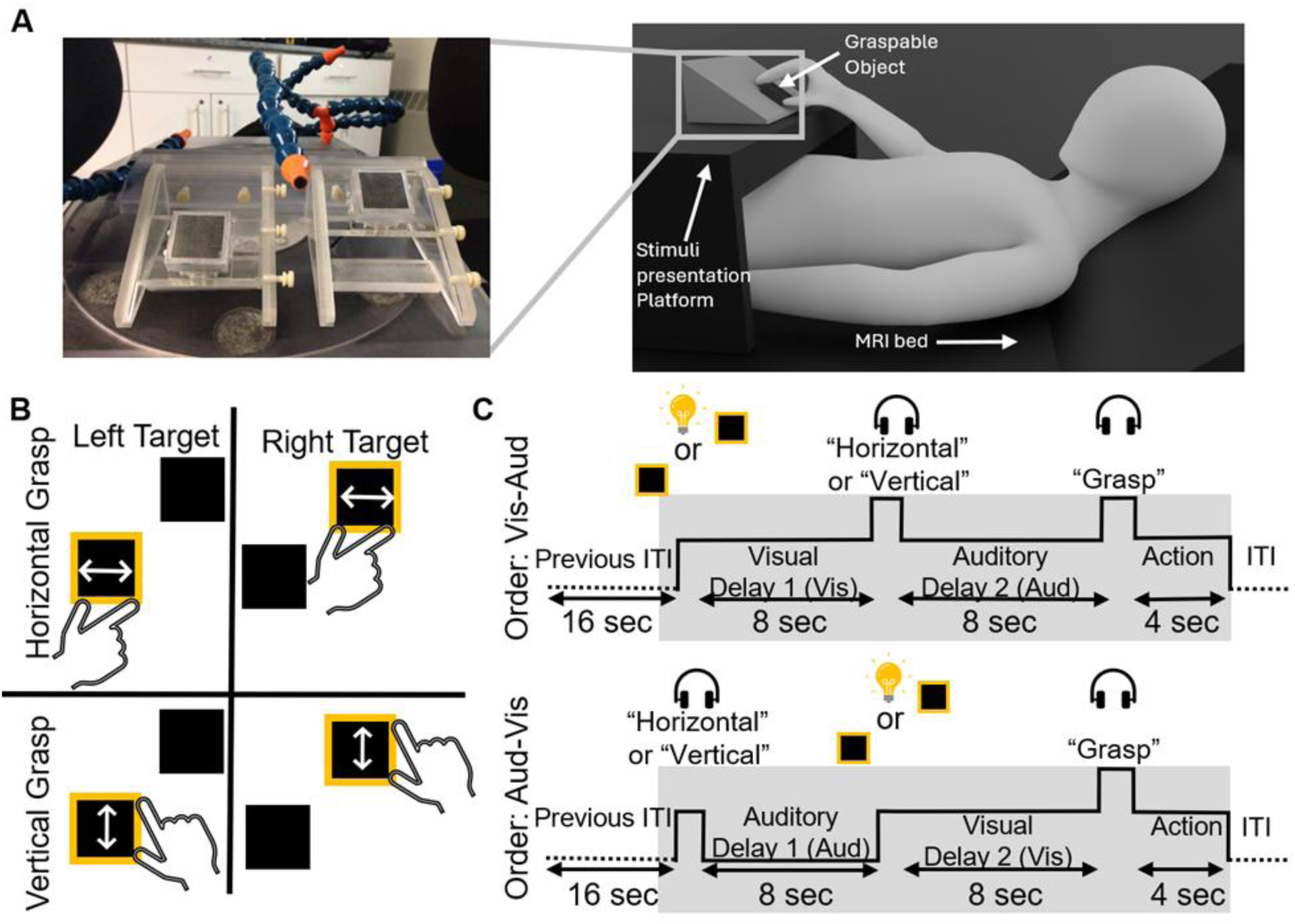
A) Experimental setup. Participants were instructed to maintain fixation on the LED light in the center, and they could grasp the object to the left or right. B) Presentation of the four possible orientations and location combinations for the grasp. C) Timeline of a single trial for each of the two trial types (Visual-Auditory and Auditory-Visual). For the Visual-Auditory trials, participants were first visually cued about the reach location, followed by an 8-second delay (Vis-Aud-D1). Then, participants were given auditory instructions about the grasp orientation, which was also followed by an 8-second delay (Vis-Aud-D2). For Auditory-Visual trials, the instructions were given in the reverse order: participants were first given auditory instructions on the grasp orientation. After an 8-second delay (Vis-Aud-D1), participants were then visually cued about the grasp location. This was followed by another 8-second delay (Aud-Vis-D2). Then, participants had 4 seconds to reach and grasp the object. Note that the auditory cues were transient and that the visual cues were sustained until the end of the task, such that participants performed a visually guided action. The inter-trial time was 16 seconds.

During experiments, two grasp stimuli were fixed to an inclined platform placed above the participants’ pelvis (Fig. 1A). Grasp stimuli consisted of two square-shaped (5 cm x 5 cm x 1.8 cm) plexiglass objects fixed to the left and right sides of the platform, at a relative height / distance chosen to optimize the biomechanical comfort of forearm motion. The objects were illuminated via underlying optic fibers to indicate which object needed to be grasped. The specific platform distance and height were adjusted for each participant so that they could comfortably reach both objects. Object orientation was fixed so that objects could be grasped with either a horizontal or vertical grip. An adjustable ‘snake’ was used to place a dim, fibre-controlled LED between the two objects for visual fixation.

All lights and audio were controlled by a custom program on a laptop PC that received signals from the MRI scanner at the start of each trial. The windows in the scanner room were blocked with black curtains, and the room lights remained off so that, except for the fixation light, nothing on the platform was visible to the participant when the object illuminator was off. Outside the scanner room, we monitored the participant’s compliance with the task using an infrared camera (MRC 12M) to record hand movements and an eye tracker (iViewX with an MRI-compatible Avotec Silent Vision system).

### 2.3. Experimental Task Design

To investigate the brain areas involved in integrating reach location and grasp orientation, we employed a cue-separation paradigm that temporally separates auditory and visual cue presentations (Baumann et al., 2009; Cappadocia et al., 2016; Chen et al., 2014). At the start of each trial (Fig. 1B), the central fixation light was illuminated, and participants maintained their gaze on it. Participants were then provided with a visual cue of the object’s location to the left or right (Visual Cue: Vis). A verbal instruction (’Horizontal’ or ‘Vertical’) telling them to grasp the object with a horizontal or vertical grip posture (Auditory Cue: Aud).

The order of these two cues (Vis-Aud vs. Aud-Vis) varied randomly, with an 8-second delay period (D1) after the first cue presentation, a second 8-second delay period (D2) after the second cue presentation and a final 4-second period to grasp the object (Fig. 1C). After each trial, participants were required to return to the starting position on the pelvis and remain still during a 16-second intertrial interval before the subsequent trial. Therefore, a complete trial was 36 seconds long.

Overall, this paradigm formed a 2 (location instruction: left, right) × 2 (orientation instruction: horizontal, vertical) × 2 (order: Vis-Aud, Aud-Vis) study, resulting in 8 conditions. Each run consisted of 16 trials; therefore, each experimental condition was repeated twice in a random order. A baseline of 8 seconds was added to the end of each run, yielding a run time of 9.73 minutes. Each participant completed 8 runs presented in a randomized order and repeated each experimental condition 16 times. Additionally, an anatomical scan for each subject was collected halfway through the data collection session. Combined with setup time, practice trials to become familiar with the task, and brief breaks between runs, an entire session lasted approximately 2.5 hours.

### 2.4. Imaging Parameters

The T2*-weighted functional experimental data were recorded using a single-shot gradient echo-planar imaging (including repetition time (TR) of 2 seconds, echo time (TE) of 30 ms, and flip angle of 90° in a 192 mm field of view (FOV), a 64 x 64 matrix size, yielding a resolution of 3.0 mm isovoxel and a 3.5 mm slice thickness with no gap). Each volume contained 35 slices, which were sampled in an ascending and interleaved order. With the functional data, a T1-weighted anatomical scan was acquired in the same orientation as the functional imaging data using a 3D acquisition sequence (192 x 256 x 256 mm FOV with repeated matrix size yielding a resolution of 1.0-mm isovoxel, inversion time of 900 ms, TR of 1900 ms, TE of 2.52 ms, and flip angle of 9°).

### 2.5. Preprocessing

Data were cleaned and analyzed using the FMRIB Software Library 6.0.7.17 (Jenkinson et al. 2012) before moving to RStudio 2024.12.1 for Region of Interest Analysis and MATLAB R2024b for Graph Theory Analysis. The first two volumes of each scan were omitted to avoid T1 saturation effects. We used high-pass temporal filtering to remove low-frequency artifacts, applied spatial smoothing with a kernel size of 6.0 mm, and performed a slice scan time correction. We motion-corrected each functional run using MCFLIRT, which applied a rigid-body transformation. Runs that showed head displacements over 1 mm were excluded from further analysis. However, one participant displayed a large head motion in the first trial during two runs; therefore, only the initial trial was excluded. Functional acquisitions were coregistered to anatomical images using the Brain-Boundary Registration in the Montreal Neurological Institute (MNI152) 1 mm standard space (Mazziotta et al. 1995). Participants’ hand movements and gaze fixations were assessed offline to confirm their compliance with the instructions. Combining all exclusion criteria across participants, fewer than 5% of trials needed to be removed from further analysis.

### 2.6. Graph Theoretical Analysis

For our functional connectivity analysis, we created 200 spheres of 3 mm radius on FSLeyes using the 200 nodes’ coordinates from the Schaefer-200 Atlas (Schaefer et al., 2018; Fig. 2A). The raw BOLD signals from each sphere for each run across all participants included in our analysis were extracted to conduct the functional connectivity analysis (Fig. 2B). For these runs, we also obtained the event files, the parameter estimates and the residuals for each region of interest to compute the contrast-reduced time series of our data. Contrast-reduced time series is a partial model fit of the voxel time course that computes contrast-related signals and residual errors while removing condition-irrelevant regressors, following the implementation described in FSL FEAT Mode (McCarthy 2025). Thus, yielding a cleaner signal that reflects condition-specific neural dynamics (Musa et al. 2025). Data from all the trials used in our analysis were extracted from the run prior to further analysis (Fig. 2C).

**Figure 2.**
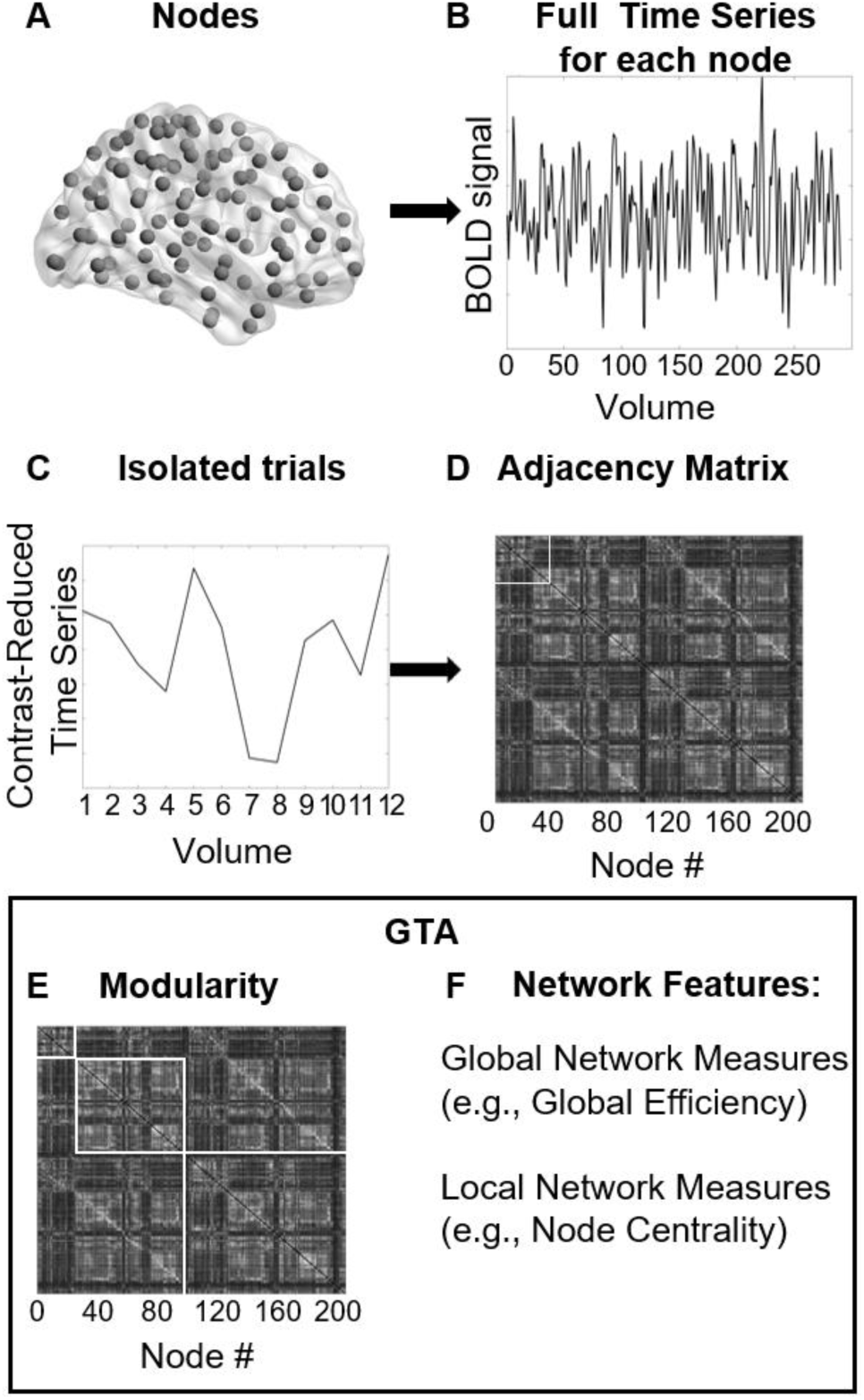
Schematic representation of graph theoretical analysis (GTA) applied to fMRI data. A) A cortical surface model illustrating the brain, with 200 regions of interest (ROIs) / nodes used for network analysis. B) The blood-oxygen-level-dependent (BOLD) signal extracted from the entire run for each ROI. C) Isolation of the contrast-reduced time series for individual trials. D) Construction of an adjacency matrix representing functional connectivity between brain nodes. E) Application of GTA to examine network properties, such as modularity, where each square delimits a module. F) Extraction of global and local network measures, including global efficiency and node centrality, to quantify brain network organization.

To construct functional networks, we calculated correlations of the contrast-reduced timeseries obtained across the 200 nodes for each trial (Musa et al. 2025; Tomou et al. 2025). Our data points started with the last two volumes of the intertrial period preceding the first cue presentation and continued until the end of the grasp period (grey box in Fig. 1C). The absolute values of the correlation outputs were placed in 200 x 200 adjacency matrices (Fig. 2D). These matrices were averaged across all trials, so that each participant had one matrix per condition. Then, the graph theoretical measures were calculated for each participant (i.e., for each matrix), and statistical comparisons were performed between conditions. However, for modularity analysis, averaging across participants yielded a single matrix per condition, which was then used to identify modules.

Then, GTA measurements were computed on these matrices using the Brain Connectivity Toolbox (Rubinov and Sporns 2010). In addition to self-developed functions for the energy and modularity score (Ghaderi et al. 2020; Ghaderi et al. 2025). Definitions of the GTA measures used are provided in Table 1. First, we divided our whole-brain network from the grand average into modules, smaller dense communities of non-overlapping nodes, which were identified by maximizing Newman-Girvan modularity for undirected networks. We maximized the number of communications within the subnetwork and minimized the number of connections (edges) across subnetworks. All brain networks were visualized using BrainNet Viewer (Xia et al. 2013). Second, we computed Newman’s *Modularity* Algorithm (Q), a measure of the quality of the division into subcommunities, by comparing the number of within-community edges to the expected number of edges in a random subnetwork of the same number of nodes (Newman 2006; Ghaderi et al. 2025). More details on the approach used can be found in Tomou and colleagues (2025).

**Table 1.**
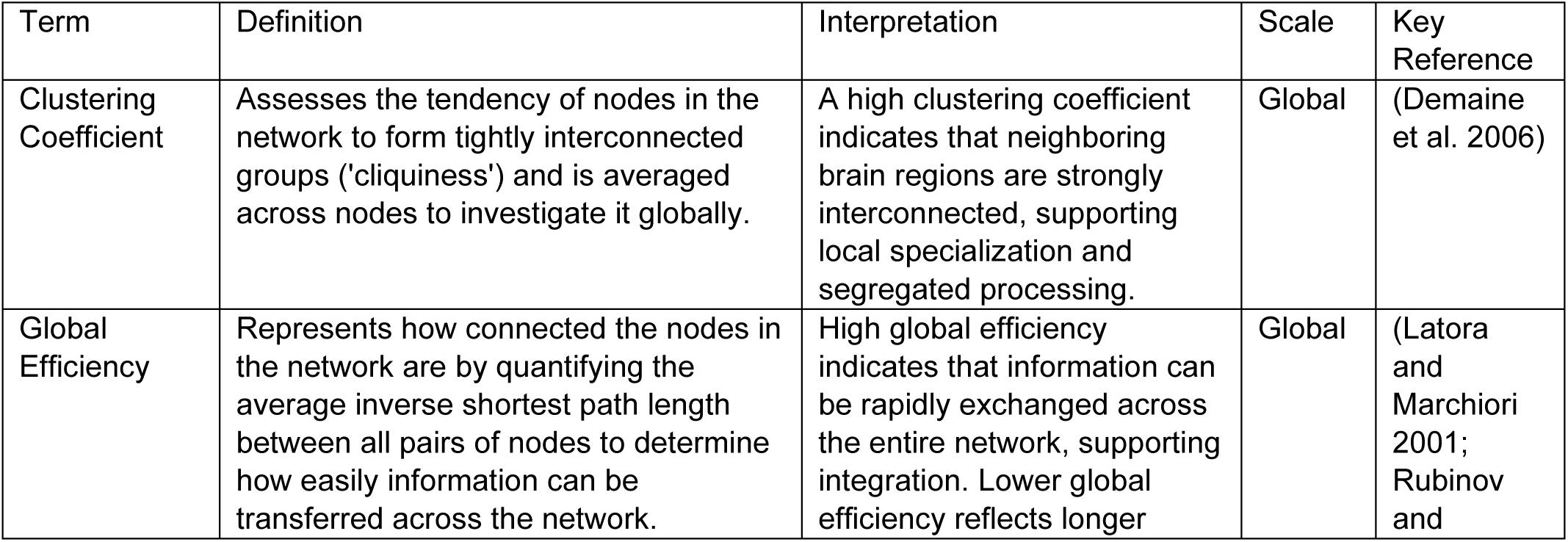
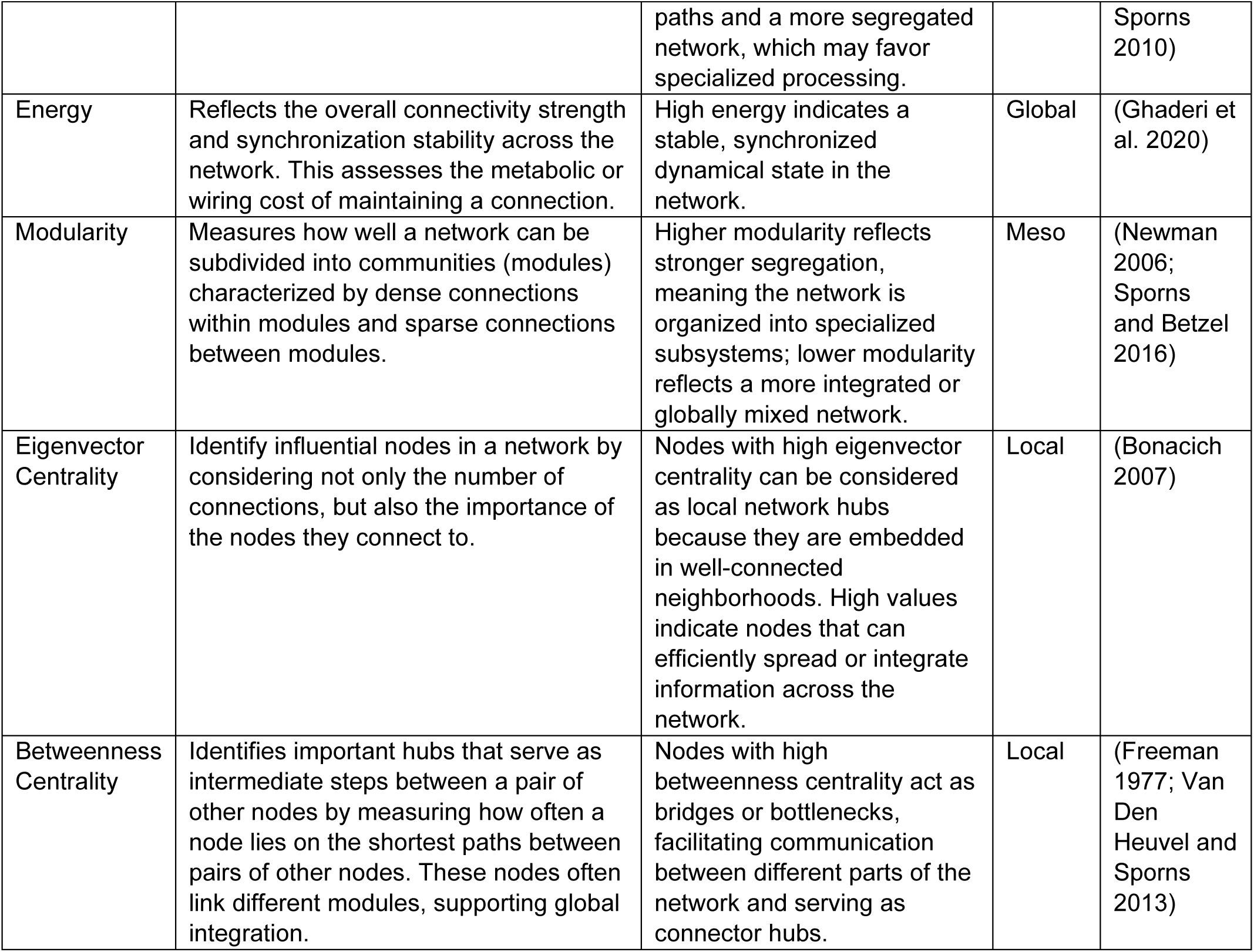
Glossary of Graph Theory Measures used for our weighted networks.

To better understand how information flows through the brain and which nodes are influential for the functional connectivity, we computed the eigenvector centrality and the betweenness centrality. The first centrality measure identified the major *eigenvector hubs,* which are well-connected nodes with high functional connectivity to other highly connected nodes, and the second centrality measure highlighted the major betw*eenness hubs,* which are regions that act as bridges between separate parts of the network (such as two different modules). These values were selected by identifying the nodes that represented the top 1% and the top 5% of the measured centrality values (Tomou et al. 2025). To capture core aspects of brain network organization, we computed the clustering coefficient, a measure of the functional segregation of the brain as the tendency of neighbour nodes to be strongly connected, the global efficiency, a measure of the functional integration which represents how efficiently information is exchanged across the whole network, and finally the energy, a measure of the dynamics and stability of synchronization in the network. All of these measures were normalized to a null model (Váša and Mišić 2022). We create these randomized datasets using a self-developed MATLAB code available for open access at https://github.com/AHGhaderi/Null-model. We computed the GTA measures using all trials combined, regardless of trial type, to assess the overall results. They were also computed separately for each trial condition (Vis-Aud or Aud-Vis) to compare differences across trial types.

Finally, we employed a machine learning approach to assess how well these graph theory features can dissociate between the two orders of cue presentation. Therefore, a supervised classification analysis was conducted using MATLAB’s *Classifier Learner* app based on each trial for all the participants. We then used the Kruskal Wallis algorithm to rank the importance of each input in their attempt to distinguish between our two conditions.

### 2.7. Statistical Analysis

*Univariate Data Analysis:* Functional MRI data were analyzed using FSL FEAT (version 6.00). At the individual run level, we used a voxelwise general linear model (GLM). Each predictor was derived from a boxcar wave function convolved with FSL’s canonical double-gamma hemodynamic response function, with temporal derivatives included to account for variability in response timing. We defined the delay phases and the action as the period of interest. We modelled our predictors using these three time periods (Delay 1, Delay 2, and Action) based on cue presentation (Vis-Aud or Aud-Vis), resulting in a total of 6 predictors of interest: Delay 1 Vis-Aud, Delay 1 Aud-Vis, Delay 2 Vis-Aud, Delay 2 Aud-Vis, Grasp Vis-Aud, and Grasp Aud-Vis. We combined all conditions (left/right and clockwise/counterclockwise) for each order of cue presentation within each period and compared them to baseline to investigate their general response to the task. For each subject, their run-level results were combined using a fixed-effects model. Finally, group-level analyses were performed using a mixed-effects model (FLAME 1), with statistical maps cluster-corrected using a z-normalized threshold of 2.3 and a corrected cluster significance threshold of P = 0.05 (Worsley 2001). The threshold was selected in accordance with previous action-related fMRI studies (Marangon et al. 2016; Gertz et al. 2017). Surface plots were generated using Nilearn functions in Python (Abraham et al. 2014).

*Region of Interest Analysis:* We hypothesized that cortical areas known to be involved in motor control would exhibit a specific increase in activity and carry information regarding the order of cue presentation. For this effort, we extracted beta weights from 7 regions of interest (V1, SPOC, pIPS, aIPS, PMv, PMd, and M1), and their values were averaged from a 6 mm radius sphere created around centroid coordinates. For V1, the coordinates were identified by converting Talairach values reported in a visually guided grasping task (Monaco et al. 2024) to MNI coordinates using seed-based d mapping according to Lancaster and colleagues (2007). In contrast, the other regions were identified from Neurosynth data terms based on meta-analyses using the terms: “reaching” and “grasping” (Yarkoni et al. 2011). The ROIs were then used to extract β-weight values for each participant in the Vis-Aud and Aud-Vis conditions during the three phases (Delay 1, Delay 2, and Action). We computed paired t-tests on their β-weights to identify regions that differ significantly from zero, and compared them across conditions (Vis-Aud vs Aud-Vis). We use the false-discovery rate based on the number of ROIs assessed to correct for multiple comparisons. Following the approach reported by Baltaretu and colleagues (2020), we conducted a post hoc power analysis to verify that our sample was sufficient to detect a difference in the order of cue presentation. Using the β-weight, which shows the smallest difference between the two conditions (at pIPS) in delay 1, we found that at our sample size, the region has an effect size (d(repeated) = 1.05) and provides power = 0.91.

*GTA analysis:* Using a community-assignment algorithm, all nodes in our network were grouped into modules. To confirm that modules exceeded the modularity expected under a null model, they were normalized to the modularity of a randomized subnetwork. A modularity score greater than 1 indicates that the connectivity within the selected group of nodes (subnetwork) is more organized than would be expected by chance. Because the normalized modularity score is strictly positive and interpreted as a relative measure, it was log-transformed, and a non-parametric one-sample sign-flip permutation test (right-tailed with 50000 permutations) was applied to statistically confirm that the results are not due to a random network. Modules with non-significant modularity scores were also reported for completeness. In addition, nonparametric permutation t-tests with 10000 randomizations were computed to compare graph theoretic features (Modularity, Cluster Coefficient, Global Efficiency, and Energy) between the two orders of cue presentation (Tomou et al. 2025). A false discovery rate (FDR: q) was calculated to obtain an adjusted p-value, controlling for multiple comparisons. Furthermore, using nonparametric permutation testing allowed us to account for the small sample size, as it avoids distributional assumptions required in parametric tests and creates a randomized dataset by reassigning condition labels. We can then assess whether our original data differs significantly from the null distribution. For our classification, we employed a 10-fold cross-validation procedure to assess the model’s performance and prevent overfitting. This approach validated the stability of our classification by dividing the data into 10 groups (folds), with the model being trained on 9 folds and tested on the remaining fold. This was repeated for 10 iterations, so that each fold served as the test fold at least once.

## 3. RESULTS

The goal of this study was to employ GTA network analysis to determine how our sensory cues (i.e., visual for object, auditory for grip strategy) were integrated for action. To justify this analysis, we first used univariate voxelwise and ROI analyses to confirm that our task produced the expected order-dependent sensory activation during cue presentation, followed by robust parietofrontal activation during reach planning and action.

### 3.1. Whole-Brain Univariate Analyses

For this analysis, we examined activation during Delay 1, Delay 2, and Action versus the baseline. Based on previous neuroimaging studies (i.e., Straube et al., 2017; Nelson and Mayhew, 2025), one might expect BOLD activation in the two delay periods to be dominated by the type and order of the sensory stimulus, i.e., visual followed by auditory cortex activation in the Vis-Aud order, and the reverse in the Aud-Vis order. During the Action phase, we expected additional activation of the fronto-parietal cortex and other areas associated with reach execution. We first computed the order-combined increase in activity relative to baseline (Vis-Aud + Aud-Vis > baseline) across the three epochs (Fig. 3, *Left Column*). During Delay 1 (Fig. 3A), we observed an increase in activity in the bilateral Early Visual Cortex (z = 5.85), the left Supramarginal Gyrus (z = 3.35), the left Precentral Gyrus (z = 3.24), and the left Postcentral Gyrus (z = 4.0). Delay 2 (Fig. 3B) showed a greater activation in the same regions: the bilateral Early Visual Cortex (z = 5.47), the left Supramarginal Gyrus (z = 3.52), the left Precentral Gyrus (z = 2.95), the left Postcentral Gyrus (z = 3.19), as well as in the Superior Temporal Gyrus (auditory cortex: A1; z = 3.98) and the Lateral Occipital Cortex (z = 2.94).

**Figure 3.**
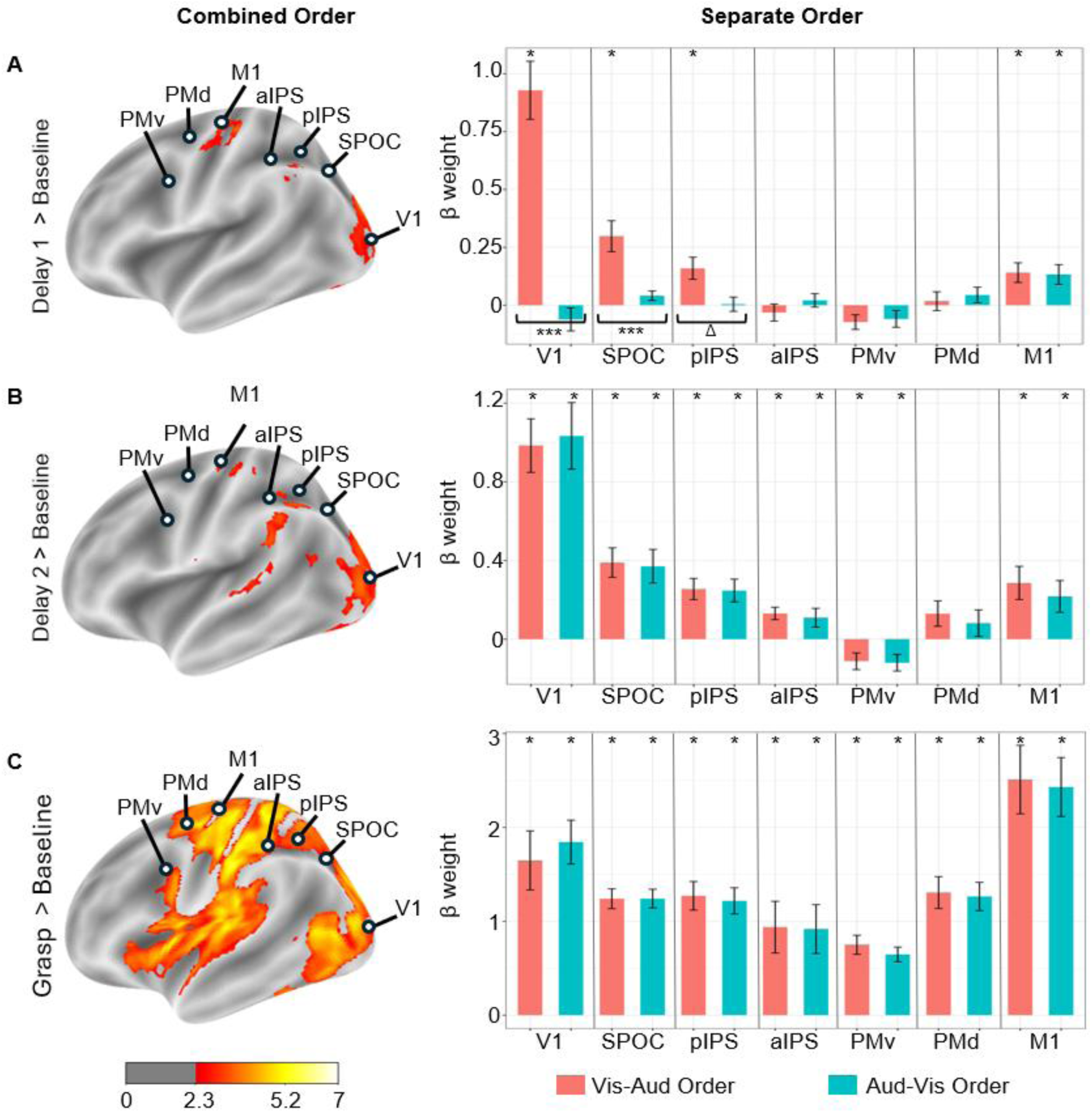
Left panel presents the lateral views of the cortical activation maps in the left hemisphere for the combined order during the three epochs, with colour bars indicating statistical intensity (z threshold = 2.3). Highlighted in circles are seven regions of interest used to plot the bar graphs. Right panel presents the average β weight (± standard error) for Visual Cortex (V1), Superior Parietal Occipital Cortex (SPOC), posterior Intraparietal Sulcus (pIPS), anterior Intraparietal Sulcus (aIPS), ventral Premotor Cortex (PMv), dorsal Premotor Cortex (PMd), Primary Motor Cortex (M1) for the Visual-Auditory Order (in pink) and the Auditory-Visual Order (in blue). Asterisks above the bars indicate regions with β weights significantly different from zero. Asterisks below the graphs indicate that the β weights differ significantly by cue presentation order, and Δ shows that the β weights differ significantly by order, but this difference did not survive correction for multiple comparisons. FDR was applied to correct for multiple comparisons. A) shows the results during the first delay period, B) shows the results during the second delay period, whereas C) shows the results during the grasp/action phase.

During the execution phase (Fig. 3C), both tasks resulted in widespread activity in the insula, the middle and posterior cingulate cortex, the superior temporal gyrus, and the striate cortex, pre- and postcentral gyrus, with highest increase in activation in visually-guided action such as the bilateral Lingual Gyrus (z = 6.1), Cuneus (z = 6.0), Lateral Occipital Cortex (z = 6.1), Superior Parietal Lobule (z = 5.87), Supramarginal Gyrus (z = 5.8), Precentral gyrus (z = 6.0), Postcentral gyrus (z = 5.8). In addition to continued activation (or reactivation) of visual and auditory sensory areas, we observed a broad swath of activation spanning all four cortical lobes during the Action Phase, including most of the well-known reach and grasp-related brain areas (Cavina-Pratesi et al. 2010; Turella and Lingnau 2014; Vingerhoets 2014).

### 3.2. Region of Interest Analysis: Visual and Motor Components of the Reach Network

To test the influence of cue order on brain activation within regions of interest (ROIs), we isolated the β weights from seven reach and grasp-related brain areas in visual and motor cortex (V1, SPOC, pIPS, aIPS, PMv, PMd, and M1 in the left hemisphere; see Table 2). To complement whole-brain analyses, we performed ROI-based analyses to test a priori hypotheses in regions implicated with motor control. Their event-related responses were evaluated relative to zero, corresponding to the implicit baseline in the inter-trial interval, before extending our analysis to examine the effect of the order of cue presentation. During Delay 1, we found a significant increase in β values for V1, SPOC, pIPS, and M1 in the Vis-Aud Order as well as M1 in the Aud-Vis Order. During Delay 2, we found a significant increase in β weights in the two orders for V1, SPOC, pIPS, aIPS, and M1, whereas PMv showed a decrease in β weight. Finally, during the grasp, we observed increased activation across all regions investigated: V1, SPOC, pIPS, aIPS, PMv, PMd, and M1. When comparing the two orders, we only found a significant difference during Delay 1 in V1 (q = 8.65 × 10^-5^), SPOC (q = 8.017×10^-3^), and pIPS (p = 0.026); however, the difference in pIPS did not survive the correction for multiple comparisons (q = 6.085×10^-2^). Overall, these findings suggest a decrease in cue-order-dependence and an increase in overall brain activity as cue information accumulated toward the final grasp performance.

**Table 2.** Summary of Means, Standard Errors, and FDR-corrected q-values for the significant difference in β weight.

|  | ROI | Order | Mean | SE | FDR q-value |
| --- | --- | --- | --- | --- | --- |
| Delay 1 | V1 | Vis-Aud | 0.929 | 0.126 | $1.905 \times 10^{-4}$ |
| | SPOC | Vis-Aud | 0.289 | 0.067 | $6.808 \times 10^{-3}$ |
|  | pIPS | Vis-Aud | 0.160 | 0.047 | 0.024 |
|  | M1 | Vis-Aud | 0.140 | 0.042 | 0.024 |
|  |  | Aud-Vis | 0.132 | 0.043 | 0.029 |
| Delay 2 | V1 | Vis-Aud | 0.984 | 0.136 | $2.375 \times 10^{-4}$ |
| | | Aud-Vis | 1.033 | 0.170 | $5.360 \times 10^{-4}$ |
| | SPOC | Vis-Aud | 0.390 | 0.075 | $1.367 \times 10^{-3}$ |
| | | Aud-Vis | 0.371 | 0.086 | $2.181 \times 10^{-3}$ |
| | pIPS | Vis-Aud | 0.255 | 0.054 | $2.181 \times 10^{-3}$ |
| | | Aud-Vis | 0.248 | 0.058 | $2.983 \times 10^{-3}$ |
| | aIPS | Vis-Aud | 0.131 | 0.032 | $3.172 \times 10^{-3}$ |
|  |  | Aud-Vis | 0.109 | 0.048 | 0.050 |
|  | PMv | Vis-Aud | -0.112 | 0.042 | 0.029 |
|  |  | Aud-Vis | -0.120 | 0.042 | 0.026 |
|  | M1 | Vis-Aud | 0.286 | 0.084 | 0.010 |
|  |  | Aud-Vis | 0.218 | 0.081 | 0.029 |
| Grasp | V1 | Vis-Aud | 1.649 | 0.315 | $3.242 \times 10^{-4}$ |
| | | Aud-Vis | 1.843 | 0.234 | $1.432 \times 10^{-5}$ |
| | SPOC | Vis-Aud | 1.240 | 0.105 | $1.013 \times 10^{-6}$ |
| | | Aud-Vis | 1.242 | 0.099 | $1.013 \times 10^{-6}$ |
| | pIPS | Vis-Aud | 1.217 | 0.152 | $1.090 \times 10^{-5}$ |
| | | Aud-Vis | 1.218 | 0.140 | $1.090 \times 10^{-5}$ |
| | aIPS | Vis-Aud | 0.938 | 0.275 | $5.881 \times 10^{-3}$ |
| | | Aud-Vis | 0.917 | 0.260 | $5.131 \times 10^{-3}$ |
| | PMv | Vis-Aud | 0.750 | 0.101 | $1.876 \times 10^{-5}$ |
| | | Aud-Vis | 0.647 | 0.078 | $1.090 \times 10^{-5}$ |
| | PMd | Vis-Aud | 1.306 | 0.169 | $1.432 \times 10^{-5}$ |
| | | Aud-Vis | 1.264 | 0.150 | $1.090 \times 10^{-5}$ |
| | M1 | Vis-Aud | 2.509 | 0.365 | $3.375 \times 10^{-5}$ |
| | | Aud-Vis | 2.430 | 0.313 | $1.432 \times 10^{-5}$ |

### 3.3. Functional Network Analysis

Having confirmed that our task generally produced the patterns of sensory and motor activation that one would expect from the literature, we proceeded to our main questions: how are our two cues (visual for object properties, and auditory for the grip plan) integrated into the cortical reach network, and to what degree are the resulting functional networks dependent on the order of cue presentation? For this purpose, we performed a functional network analysis using *graph theory*, based on correlations between the contrast-reduced timeseries of our various cortical nodes (Fig. 2). In this analysis, we examined the time period from the end of the previous intertrial period of each trial up to the end of the grasp period, both for the combined and separate cue orders.

#### 3.3.1. Global Network Properties

We computed the global network features (clustering coefficient, global efficiency, energy) normalized with a null distribution to ensure that our data formed a ‘real’ (not random) network and compared their values for the two cue presentation orders (Fig. 4). A clustering coefficient value equal to or above 1 represents a non-random network, which is evident in the combined clustering coefficient (mean = 1.002, SE = 1.855×10^-4^, p = 9.998×10^-5^) as well as the separate order clustering coefficient (Vis-Aud: mean = 1.002, SE = 1.765×10^-4^, p = 2.00×10^-4^ and Aud-Vis: mean = 1.002, SE = 2.011×10^-4^, p = 2.40×10^-4^; Fig. 4 A-B). A global efficiency value smaller than 1 represents a non-random network, as observed in both the combined global efficiency of the combined condition (mean = 0.9918, SE = 0.0012, p = 3.199×10^-4^) and the separate order global efficiency (Vis-Aud: mean = 0.990, SE = 0.0015, p = 2.799×10^-4^ and Aud-Vis: mean = 0.9905, SE = 9.437 x 10^-4^, p = 2.20×10^-4^; Fig. 4 C-D). Similarly, an Energy value smaller than 1 represents a non-random network, as found in both the combined energy (mean = 0.8434, SE = 0.0085, p = 2.00×10^-4^) and the separate orders (Vis-Aud: mean = 0.8440, SE = 0.0075, p = 3.199×10^-4^ and Aud-Vis: mean = 0.8438, SE = 0.0084, p = 2.40×10^-4^; Fig. 4 E-F). However, our permutation t-tests comparing the global measures between the Vis-Aud and Aud-Vis conditions were not significant: clustering coefficient (t(11) = -0.188, p = 0.858), global efficiency (t(11) = -0.181, p = 0.859), and energy (t(11) = 0.044, p = 0.964). Overall, these analyses supported the notion that these are ‘real’ networks, even when separated into the two order conditions, but a more detailed analysis at the mesoscale (modular) and local (hub) layers was required to test our hypotheses.

**Figure 4.**
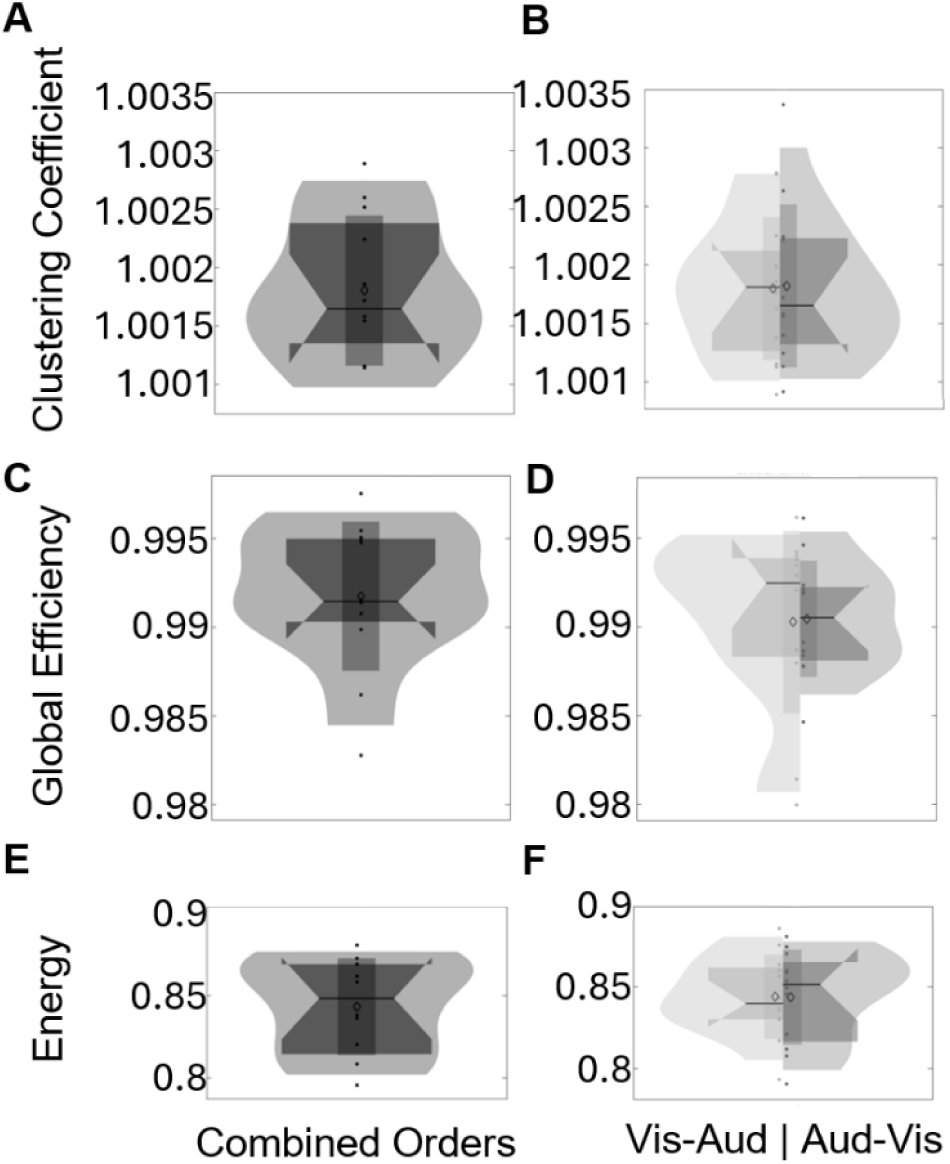
Violin plots displaying distributions of normalized global network parameters: clustering coefficient (A, B), global efficiency (C, D), and energy (E, F). The left column (A, C, E) plots the distributions of the whole dataset (combined orders). The right column (B, D, F) plots the distributions of the Vis-Aud order vs. Aud-Vis order.

#### 3.3.2. Correlated Time Series Suggest Parallel Processing Streams

Our hypothesis was that sensorimotor integration in this task would be modular, i.e., visual signals should be integrated into the motor system via the parietal cortex, whereas auditory signals could be integrated via frontal centres. If this is correct, then the BOLD time series in the visual cortex should tend to correlate with parietal signals, whereas those in the auditory cortex should correlate with frontal signals.

To illustrate this, we selected example regions from the Schaeffer atlas whose locations overlapped with areas of peak cortical activity (Fig. 5). The time series for primary visual cortex (A) resembled and significantly correlated with the time course for superior parietal lobule (B) in both presentation orders (overall r = 0.762, t(10)=3.72, q = 0.008 for Vis-Aud order and r = 0.932, t(10) = 8.13, q = 4.083×10^-5^ for Aud-Vis order). Likewise, the time course for the peak voxel within primary auditory cortex (C) resembles and significantly correlates with the time course for the peak voxel in dorsal premotor cortex (D) (overall r = 0.962, t(10) = 11.12, q = 4.763×10^-6^ for Vis-Aud order and r = 0.845, t(10) = 5.00, q = 0.003 for Aud-Vis order). In contrast, the opposite correlations (V1-premotor, A1-SPL) were lower and non-significant (r = 0.490, q = 0.141 and r = 0.619, q = 0.051 for Vis-Aud order; r = -0.025, q = 0.939 and r = 0.080, q = 0.9188 for Aud-Vis order). These specific examples appear to support our hypothesis. The next section employs GTA analysis to quantitatively test this hypothesis for 200 nodes distributed across the entire cortex.

**Figure 5.**
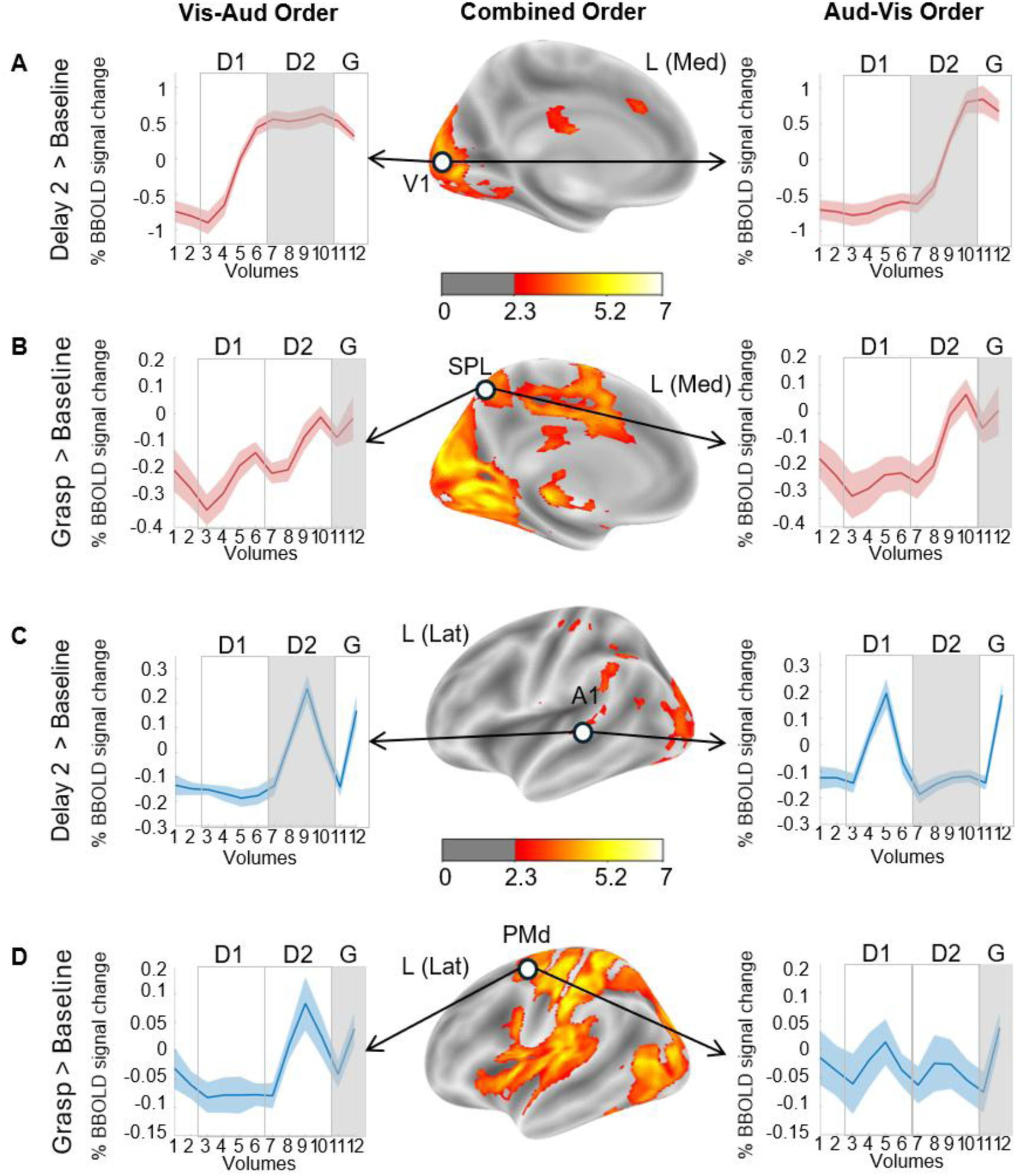
Medial and lateral views of the cortical activation maps for the Vis-Aud and Aud-Vis conditions during the Delay 2 and Grasp epochs are presented in the left hemisphere, with colour bars indicating statistical intensity (z threshold = 2.3). Highlighted in circle are two cue acquisitions regions (Primary Visual Cortex [V1; MNI: -5, -93, -4] and Primary Auditory Cortex [A1; MNI: -51, -4, -2]) and two regions involved in preparation for action (dorsal Premotor Cortex [PMd; MNI: -43, 6, 43], Superior Parietal Lobule [SPL; MNI: -17, -53, 68]) from which the average percentage changed in BOLD signal were extracted (± standard error). On the left, this data is presented for the Visual-Auditory (Vis-Aud) condition; on the right, for the Auditory-Visual (Aud-Vis) condition. A & C) show the results during the second delay period; B & D) show the results during the grasp/action phase. These regions are highlighted in a grey-shaded square in the timeseries. The 12 data points represent time in terms of volumes within a trial (D1: Delay 1, D2: Delay 2, G: Grasp). Shaded regions indicate variability across participants. Time-course data are color-coded red for V1 (in A and B) and SPL, and blue for A1 and PMd (in C and D).

#### 3.3.3. Modularity Analysis: Occipital-Parietal vs. Temporal-Frontal Modules

A formal modularity analysis provides the means to extend and quantify the examples shown in Figure 5 by identifying clusters of highly correlated BOLD time series (Figure 2 E) between our 200 nodes (Figure 2A). Figure 6A shows the results of such an analysis in our complete, order-combined dataset. This analysis subdivided our nodes into three putative modules: a Temporal-Frontal-Somatomotor module spanning the central sulcus (Module 1, blue), another module spanning Occipital-Parietal-Somatomotor cortex (Module 2, red), and a third module (Module 3, yellow) spanning the remaining nodes. We will summarize the functional anatomy of these putative modules qualitatively and then quantify / test their significance in the following paragraphs.

**Figure 6.**
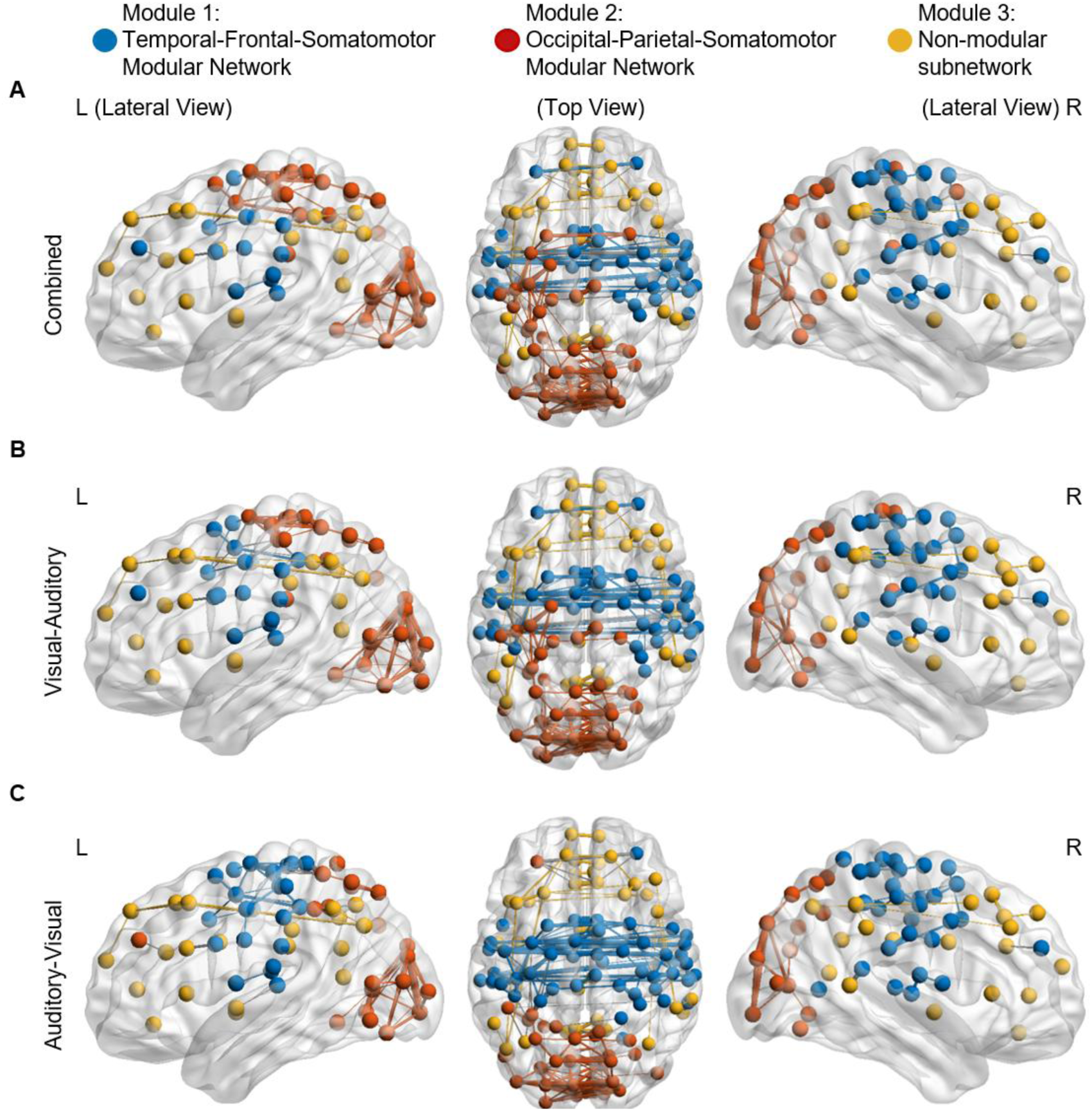
Network modularity analysis for the Visual-Auditory and Auditory-Visual conditions. A) Combined network modularity across both conditions. B) Modular structure of functional brain networks during the Visual-Auditory condition. B) Modular structure during the Auditory-Visual condition. Nodes represent brain regions, and edges represent the functional connections between them. Modules are color-coded: the Occipital-Parietal-Somatomotor Module (red), the Temporal-Frontal-Somatomotor Module (blue), and the non-significant module (yellow). Three views of the brain are shown: right hemisphere (R), top-down, and left hemisphere (L). This analysis highlights distinct modular organization patterns associated with each condition.

**Module 1** (blue) was largely bilateral and appears to be involved in relaying the auditory instruction for the grip strategy. This module included areas in the Superior Temporal Gyrus that were activated during auditory cue presentation, as well as regions involved in preparation and execution of action, including Lateral Prefrontal Cortex, Dorsal and Ventral Premotor Cortex, and most portions of primary somatomotor cortex (preparation and execution of action; Rizzolatti and Luppino, 2001; Monaco et al., 2024).

**Module 2** (red) appears to relay visual information about the stimulus to the motor system. This module included bilateral occipital areas (such as the Striate Cortex and Extrastriate Cortex) that were activated by the visual stimulus during the Delay periods, and bilateral posterior parietal areas (including the Superior Parietal Lobule and Intraparietal Sulcus) implicated in visually guided reach and grasp (Grefkes and Fink 2005; Buneo and Andersen 2006; Culham and Valyear 2006; Gallivan and Culham 2015). This module also ‘invaded’ portions of the left somatomotor cortex that were likely related to right-hand motion (Penfield and Boldrey 1937; Kim et al. 1993).

**Module 3** (yellow) spans prefrontal cortex and portions of inferior parietal and temporal cortex, however, this module did not reach significance.

**Order-Dependence.** When the data were divided by order (Fig. 6B-C), the same basic modular organization persisted (i.e., a Temporal-Frontal-Somatomotor module and an Occipital-Parietal-Somatomotor module), but with some subtle differences. For example, when the visual cue was presented first (Fig. 6B), there was a continued invasion of connectivity of the Occipital-Parietal-Somatomotor module into the left somatomotor cortex. In contrast, when the orientation cue was presented first (Fig. 6C), the Occipital-Parietal-Somatomotor module was more restricted, and the blue module fully spanned both sides of the Somatomotor cortex, decreasing lateralization in both modules.

Overall, these results suggest that 1) information from the visual cue dominated the ‘dorsal stream’ visual areas involved in visually guided movement, 2) the auditory cue (for the grip plan) was incorporated into more frontal mechanisms, and 3) these modules were largely independent of cue order, except that the initial cue (auditory or visual) tended to dominate signals in the hand-related portion of left somatomotor cortex.

To quantify these results, we computed the modularity score (Q) for each sub-network, first in the combined dataset (Fig. 7A). Out of the three communities, modularity was significantly higher than the null model for two modules: Module 1 (mean log value = 0.163; q = 4.799×10^-4^) and Module 2 (mean log value = 0.195, q = 4.799×10^-4^), but not for Module 3 (mean log value = -0.032, q = 0.998). There was a significant difference (t(11) = -8.637, q = 4.693×10^-6^) in modularity between Module 1 (mean = 1.179, SE = 0.022) and Module 3 (mean = 0.9684, SE = 0.007). There was also a significant difference (t(11) = -9.280, q = 4.650×10^-6^) between Module 2 (mean = 1.218, SE = 0.024) and Module 3. However, there was no significant difference in modularity between Module 1 and Module 2 (t(11) = 0.913, q = 0.379).

**Figure 7.**
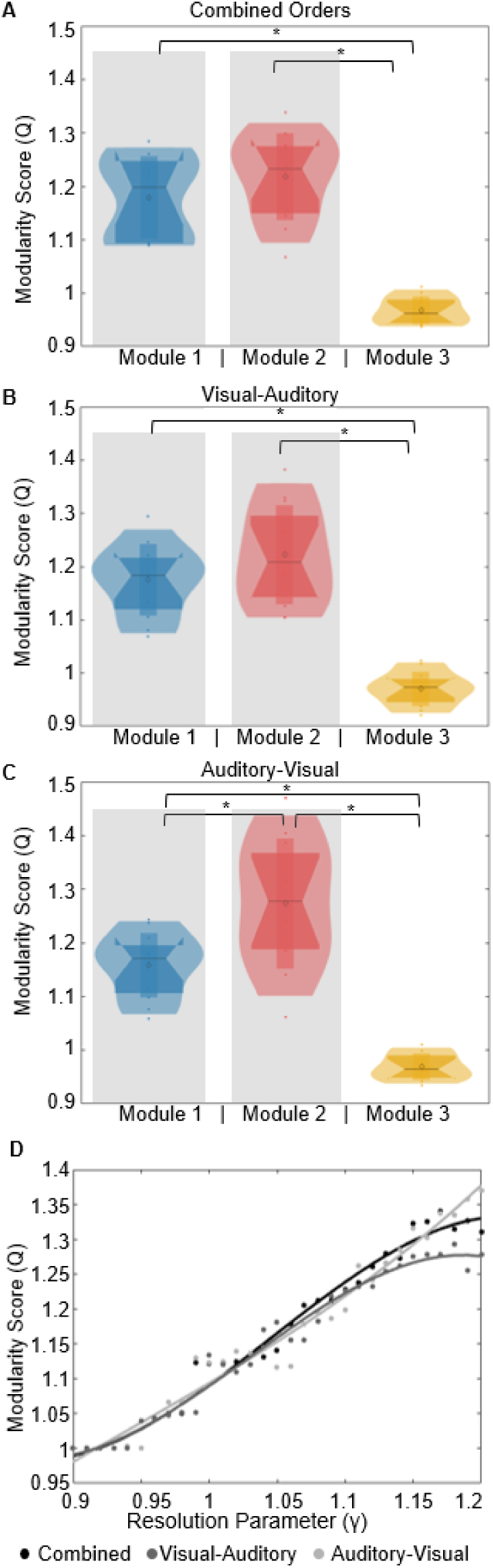
Modularity comparisons across experimental conditions and gamma values. A- C) Violin plots of modularity values for identified network modules when the orders are combined (A), in the Visual-Auditory condition (B), and the Auditory-Visual condition (C). Modules are color-coded: Module 1 (blue), Module 2 (red), and Module 3 (yellow). Asterisks (*) indicate significant differences between modules with q<0.05, and the grey boxes indicate modules with significantly greater than a random network. D) Modularity values as a function of resolution parameter for the Combined Order (black), Visual-Auditory Order (dark gray), and Auditory-Visual Order (light gray). Points represent data, and lines indicate best-fit curves. This analysis highlights differences in network modularity across task conditions and tuning parameters.

We then did the same for the order-separated networks (Fig. 7B-C). In both cases, the normalized modularity values were significantly greater than expected from a null model for Module 1 (mean log value_Vis-Aud_ = 0.160, q = 4.499×10^-4^; mean log value_Aud-Vis_ = 0.146, q = 2.999×10^-4^) and Module 2 (mean log value_Vis-Aud_ = 0.198, q = 4.999×10^-4^; mean log value_Aud-Vis_ = 0.237, q = 2.999×10^-4^). In the Vis-Aud condition, there was a significant difference between Module 1 (mean = 1.175, se = 0.019) and Module 3 (mean = 0.971, se = 0.009; t(11) = -9.421, q = 4.011×10^-6^) and between Module 2 (mean = 1.222, se = 0.027) and Module 3 (t(11) = -7.469, q = 1.872×10^-5^). There was no significant difference between Module 1 and Module 2 (t(11) = 1.163, q = 0.276). However, there was a significant difference among all three modules in the Aud-Vis order (Fig. 7C): between Module 2 (mean = 1.273, se = 0.035) and Module 3 (mean = 0.969, se = 0.007; t(11) = - 7.917, q = 1.082×10^-5^), Module 1 (mean = 1.158, se = 0.017) and Module 2 (t(11) = 2.333, q = 0.040), Module 1 and Module 3 (t(11) = -9.590, q = 3.362×10^-6^). This pattern indicates that cue order does not induce a uniform network-wide change, but instead differentially affects subnetworks, likely reflecting their respective roles in processing recent versus previously integrated information.

Finally, there was a significant order-dependent difference in modularity for Module 2 (t(11) = -3.036, q = 0.035), but not in the other modules (t(11) = 0.143, q =0.263 for Module 1; t(11) = 0.215, q = 0.832 for Module 3). These comparisons were performed at the standard modularity resolution parameter (γ) of 1, but modularity showed fairly consistent increases in all three datasets as the γ values were changed from 0.9 to 1.2 (Fig. 7D).

#### 3.3.4. Classification Analysis: Cue Order-Dependence

The analysis provided above suggests that global network parameters did not distinguish cue order-dependence, whereas the modularity parameter was modestly successful at distinguishing order. To confirm this objectively, we applied a classifier algorithm to identify which of our GTA measurements (Modularity, Clustering Coefficient, Global Efficiency and Energy) could best distinguish between the order of cue presentation. Decoding was performed using a Narrow Neural Network classification algorithm.

The model achieved a positive predictive (PPV) of 59.8% (95% Confidence Interval (CI): 55.5% to 62.3%) with a false discovery rates (FDR) of 40.2% (95% CI: 37.7% to 44.5%) for the Vis-Aud order, and a PPV of 59.9% (95% CI: 55.9% to 63.1%) with an FDR of 40.1% (95% CI: 36.9% to 44.1%) for the Aud-Vis order. The Kruskal-Wallis feature selection analysis showed that the Modularity scores from Module 2 (Occipital-Parietal-Somatomotor Modular Network), followed by Module 1 (Temporal-Frontal-Somatomotor Modular Network), were the most meaningful in differentiating between the two orders of cue presentation. This demonstrates that although overall classification accuracy was modest, the model was able to distinguish between the orders above chance level and confirms that modularity was the most important parameter.

#### 3.3.5. Eigenvector Centrality Hubs

Finally, we consider which nodes in our network acted as important ‘hubs’. Eigenvector centrality was used to compute local (i.e., within-module) hubs. Fig. 8A shows the top 95th- and 99th-percentile eigenvector hubs in our modules when the orders are combined, as well as for the two cue presentation orders (Table 3). Hubs in Module 1 (blue) included nodes in bilateral dorsal anterior Cingulate Cortex (ROI47 & ROI146). Hubs for Module 2 (red) included Somatomotor Area (ROI78) and multiple occipital regions (in the left hemisphere – ROI37, ROI41, ROI46, ROI91, and ROI92 – and in the right hemisphere –ROI145 and ROI190). Overall, these hubs corresponded to known components of the ‘salience/attention’ network (Seeley et al. 2007; Schaefer et al. 2018; Seeley 2019) and occipito-parieto-frontal reach system.

**Figure 8.**
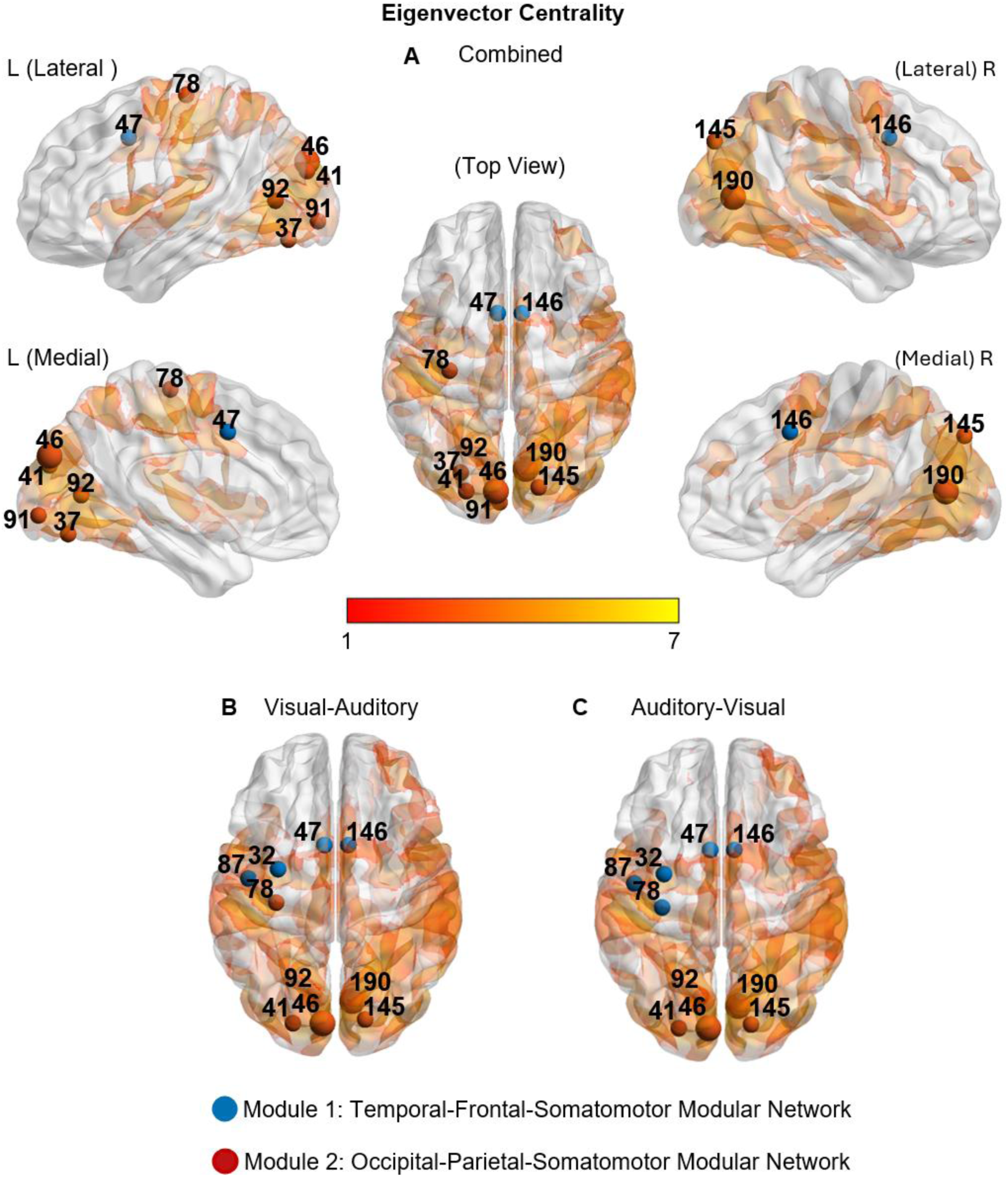
Eigenvector centrality hubs presented on brain maps overlay with activation in the action phase for (A) the combined order, (B) the Visual-Auditory Order, and (C) the Auditory-Visual Order. Smaller hubs correspond to nodes with an eigenvector centrality above 95%, and larger hubs correspond to those with an eigenvector centrality above 99%. Blue nodes are part of the Module 1 – Temporal-Frontal-Somatomotor Module, and Red nodes are part of the Module 2 – Occipital-Parietal-Somatomotor Module.

**Table 3:** Eigenvector Centrality Hubs.

| ROI # | ROI Name | Network | X | Y | Z | Module |
| --- | --- | --- | --- | --- | --- | --- |
| 32 | Left Hemisphere Frontal Eye Fields parcel 1 | Dorsal Attention Network B | -31 | -4 | 53 | 1 |
| 37 | Left Hemisphere Visual Central Extrastriate parcel 1 | Visual Central Network | -26 | -77 | -14 | 2 |
| 41 | Left Hemisphere Extrastriate parcel 5 | Visual Central Network | -23 | -87 | 23 | 2 |
| 46 | Left Hemisphere Superior Extrastriate parcel 2 | Visual Peripheral Network | -7 | -87 | 28 | 2 |
| 47 | Left Hemisphere Frontal Medial parcel 1 | Salience/Ventral Attention Network A | -6 | 9 | 41 | 1 |
| 78 | Left Hemisphere Somatomotor A parcel 4 | Somatomotor Network A | -32 | -22 | 64 | 1,2 |
| 87 | Left Hemisphere Central parcel 2 | Somatomotor Network B | -47 | -9 | 46 | 1 |
| 91 | Left Hemisphere Striate parcel 1 | Visual Central Network | -5 | -93 | -4 | 2 |
| 92 | Left Hemisphere Striate Calcarine parcel 1 | Visual Peripheral Network | -11 | -70 | 7 | 2 |
| 145 | Right Hemisphere Superior Extrastriate parcel 3 | Visual Peripheral Network | 16 | -85 | 39 | 2 |
| 146 | Right Hemisphere Frontal Medial parcel 1 | Salience/Ventral Attention Network A | 7 | 9 | 41 | 1 |
| 190 | Right Hemisphere Striate Calcarine parcel 1 | Visual Peripheral Network | 9 | -75 | 9 | 2 |

Note: Col 1: ROI Numbers, Col 3-5 MNI coordinates, Col 6: Module associated with these hubs. ROIs derived from the 200 parcellation from the Schaefer 17 Networks Atlas (Schaefer et al. 2018)

When the cue orders were separated into Vis-Aud (Fig. 8B) and Aud-Vis (Fig. 8C), there was an overall decrease in the number of hubs in Module 2 (ROI91 – Left Striate Cortex and ROI37 – Left Extrastriate) and an increase in the number of hubs in Module 1 (ROI32 – Left Frontal Eye Field and ROI87 – Left Precentral Gyrus). Comparing the two orders, ROI78 (Left Precentral Gyrus) switched between modules, tending to join the module dominated by the first sensory cue (i.e., Module 1 for Aud-Vis, Module 2 for Vis-Aud). Otherwise, the hubs for the two cue orders were nearly identical, i.e., the same ROIs were identified as important hubs, and they showed the same module assignments.

#### 3.3.6. Betweenness Centrality Hubs

Betweenness centrality hubs represent nodes with broader ‘connections’, i.e., correlations that tend to cross between modules. Fig. 9 (Table 4) shows nodes with the top 95th and 99th percentiles of betweenness centrality, color-coded by module. When the orders were combined, the majority of the hubs were part of Module 1 (blue) and were located in the bilateral Frontal regions (ROI47, ROI86, ROI146, and ROI148), the bilateral Auditory Cortex (ROI84 and ROI182), and the Parietal Operculum Cortex (ROI160). In addition, there were one hub from Module 2 (red) hub in the right Occipital Cortex (ROI137) and two hubs from Module 3 (yellow) in the right Insula (ROI192) and Frontal Operculum Cortex (ROI148).

**Figure 9.**
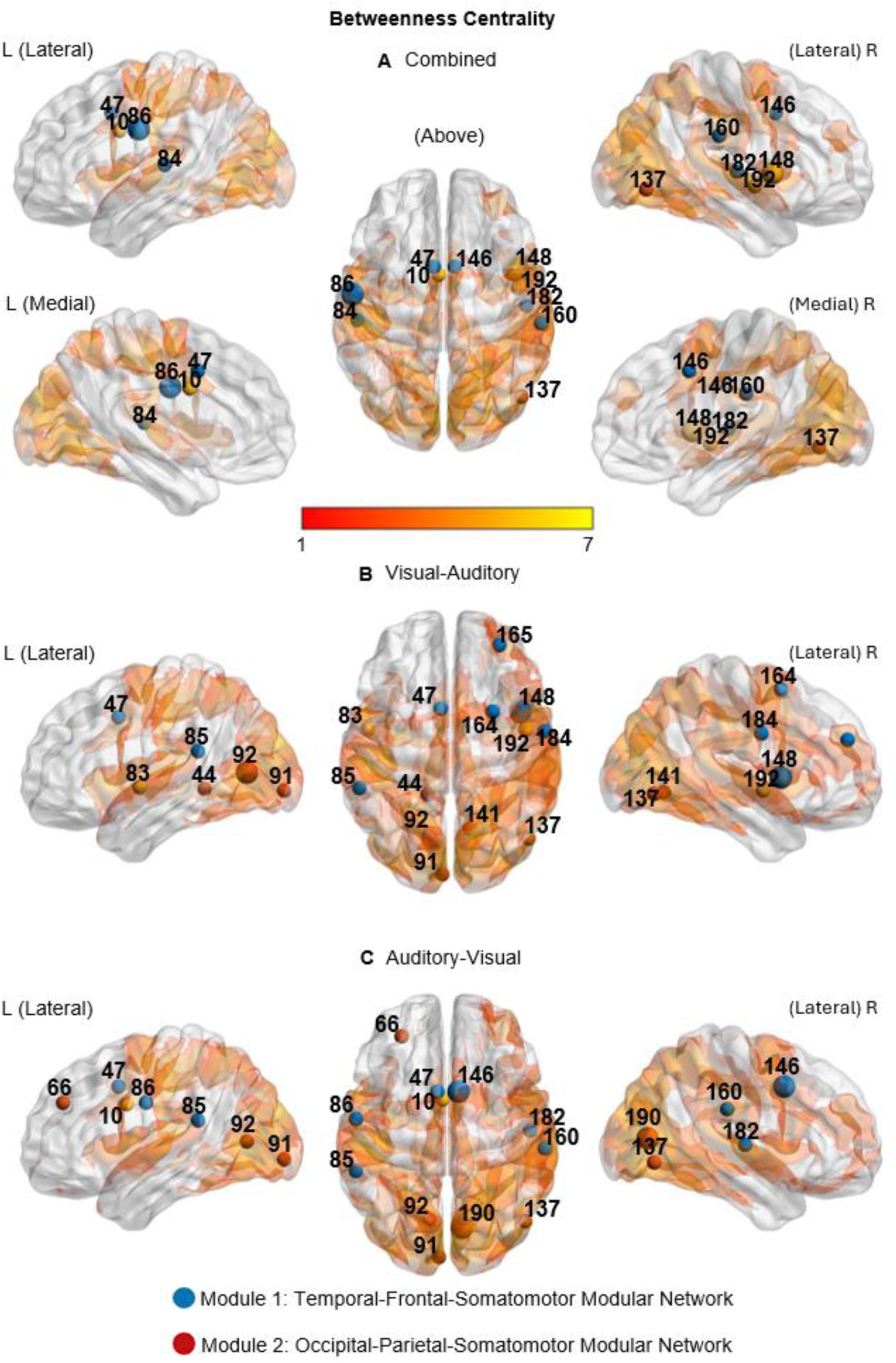
Betweenness centrality hubs are presented on brain maps overlaid with activation in the action phase for (A) the combined order, (B) the Visual-Auditory Order, and (C) the Auditory-Visual Order. Smaller hubs correspond to nodes with betweenness centrality above 95%, and larger hubs correspond to nodes with eigenvector centrality above 99%. Blue nodes are part of the Module 1 – Temporal-Frontal-Somatomotor Module. Red nodes are part of the Module 2 – Occipital-Parietal-Somatomotor Module.

**Table 4:** Betweenness Centrality Hubs.

| ROI # | Name | Network | X | Y | Z | Module |
| --- | --- | --- | --- | --- | --- | --- |
| 10 | Left Hemisphere Middle Cingulate parcel 1 | Control network A | -3 | 4 | 30 | 3 |
| 44 | Left Hemisphere Inferior Extrastriate parcel 3 | Visual Peripheral network | -14 | -44 | -3 | 2 |
| 47 | Left Hemisphere Frontal Medial parcel 1 | Saliency/Ventral Attention network A | -6 | 9 | 41 | 1 |
| 66 | Left Hemisphere Lateral Prefrontal Cortex parcel 1 | Saliency/Ventral Attention network B | -28 | 43 | 31 | 2 |
| 83 | Left Hemisphere Auditory parcel 1 | Somatomotor network B | -51 | -4 | -2 | 3 |
| 84 | Left Hemisphere Auditory parcel 2 | Somatomotor network B | -53 | -24 | 9 | 1 |
| 85 | Left Hemisphere Auditory parcel 3 | Somatomotor network B | -56 | -40 | 20 | 1 |
| 86 | Left Hemisphere Central parcel 1 | Somatomotor network B | -56 | -8 | 31 | 1 |
| 91 | Left Hemisphere Striate parcel 1 | Visual Central network | -5 | -93 | -4 | 2 |
| 92 | Left Hemisphere Striate Calcarine parcel 1 | Visual Peripheral network | -11 | -70 | 7 | 2 |
| 137 | Right Hemisphere Extrastriate parcel 2 | Visual Central network | 48 | -71 | -6 | 2 |
| 141 | Right Hemisphere Inferior Extrastriate parcel 1 | Visual Peripheral network | 12 | -65 | -5 | 2 |
| 146 | Right Hemisphere Frontal Medial parcel 1 | Saliency/Ventral Attention network A | 7 | 9 | 41 | 1 |
| 148 | Right Hemisphere Frontal Operculum parcel 1 | Saliency/Ventral Attention network A | 43 | 7 | 4 | 1 |
| 160 | Right Hemisphere Parietal Operculum parcel 1 | Saliency/Ventral Attention network A | 60 | -26 | 27 | 1 |
| 164 | Right Hemisphere Dorsal Prefrontal Cortex parcel 1 | Control network A | 26 | 7 | 58 | 1 |
| 165 | Right Hemisphere Lateral Prefrontal Cortex parcel 1 | Saliency/Ventral Attention network B | 30 | 48 | 27 | 1 |
| 182 | Right Hemisphere Auditory parcel 1 | Somatomotor network B | 51 | -15 | 5 | 1 |
| 184 | Right Hemisphere Central parcel 1 | Somatomotor network B | 58 | -5 | 31 | 1 |
| 190 | Right Hemisphere Striate Calcarine parcel 1 | Visual Peripheral network | 9 | -75 | 9 | 2 |
| 192 | Right Hemisphere Insula parcel 2 | Saliency/Ventral Attention network A | 46 | -4 | -4 | 3 |

Note: Col 1: ROI Numbers, Col 3-5 MNI coordinates, Col 6: Module associated with these hubs. ROIs derived from the 200 parcellation from the Schaefer 17 Networks Atlas (Schaefer et al. 2018)

When the two orders were investigated separately, hubness was more distributed throughout the cortex, included more occipital nodes, and showed greater variability between the two conditions (compared to Eigenvector Hubs). Specifically, the two orders have four order-independent hubs, two of which are part of Module 1 (ROI47 in the left Frontal Cortex and ROI85 in the left Auditory Cortex), and three of which are part of Module 2 (ROI91 and ROI92 in the bilateral Occipital Cortex and ROI137 in the right Occipital Cortex). In addition, the Visual-Auditory condition had distinct hubs including four regions in the right Frontal Cortex from Module 1 (ROI165, ROI148, ROI164 and ROI184), three hubs part of the Occipital Cortex in Module 2 (in the left hemisphere: ROI41, ROI91 and in the right hemisphere: ROI141) and bilateral Temporal region as part of the Module 3 (ROI83 and ROI192). As for the Auditory-Visual Order, the hubs in Module 1 also included the right Frontal region (ROI146), the left Somatomotor area (ROI86), the right Auditory Cortex (ROI182), and the right Parietal Operculum Cortex (ROI160). Module 2 included hubs from the right Occipital regions (ROI190) and the left Lateral Prefrontal Cortex (ROI66). Finally, the non-significant module included hubs in the left Cingulate Cortex (ROI10). In short, compared to eigenvector hubs, betweenness centrality hubs tended to occupy more sensory nodes and were more cue order-dependent.

## 4. Discussion

Movement can be initiated by multisensory top-down and bottom-up signals, which are integrated to achieve the desired action. In this study, we explored 1) how such cues (verbal instruction for grip orientation and visual cues for object attributes) are integrated into the motor system, and 2) whether this is influenced by the order of presentation. Overall, we found that cue integration is not undertaken by a single region or evenly distributed across all cortical areas. Instead, we found that cue-related information was processed through cortical sub-modules that progressed from the sensory cortex, communicated laterally, and finally converged in the somatomotor cortex.

Specifically, our univariate analysis revealed order-dependent responses in the sensory cortex, followed by widespread order-independent activation during the planning and execution phases. The analysis of global network properties showed that we were dealing with a non-random network with order-independent global parameters. Mesoscale analysis revealed that the data can be divided into three modules, with two significantly modular subnetworks: the temporal-frontal-somatomotor and occipital-parietal-somatomotor modules. These modules included hubs in the frontal and occipital cortex, respectively, whereas ‘betweenness’ hubs (thought to be involved in intermodular communication) were found in the auditory / frontal cortex, and the occipital cortex, respectively, with additional hubs in the non-significant module. In our classification analysis, we found that the modularity index values for these two modules were the most effective parameter for distinguishing between the order of cue presentation. We will discuss each of these results, in particular their relation to cue integration and its order-dependence, below.

### 4.1. Univariate Analysis: Activation and Order Dependence

During cue presentation phases, activation was primarily observed in sensory-related regions (Fig. 3), with a transient response in A1 during the verbal instruction and a more sustained response in V1 once the grasp target was illuminated. As expected, these responses were order-dependent, most clearly arising first in the sensory area corresponding to the modality of the first cue. In addition, we observed activation in restricted premotor and sensorimotor regions during cue presentation, consistent with advanced processing of visual and auditory information and preparation for the upcoming action (Hall et al. 2000; Vannest et al. 2009; Singhal et al. 2013).

As participants transitioned through the different phases of our task, we observed a typical increase in brain activity, culminating in widespread, order-independent cortical activation during the action execution phase (Begliomini et al. 2008; Brandi et al. 2014; Cappadocia et al. 2017; Klein et al. 2023). This included occipital-parietal-frontal areas, which are required to accomplish visually-guided movement (Vesia and Crawford 2012; Gallivan and Culham 2015). In addition, we observed activation within the ‘action mode’ network, including cingulo-opercular areas thought to support the integration of bottom-up visual information with task-relevant information (Yu et al. 2023; Dosenbach et al. 2025). These data suggest that cue integration is occurring, but do not show how.

None of these results were surprising, and indeed that was the point of this analysis: to show that our task showed the expected order-dependent sensory activation and robust sensorimotor activation required for our network analysis.

### 4.2. Graph Theory Analysis of Functional Network Properties

The main assumption behind this study is that multimodal cue integration for action is accomplished through functional network properties, rather than specific sites. The basic concept behind functional network analysis is that signal correlations reflect the propagation of information through a network (Sporns 2002). This is illustrated in Figure 5, where all four example regions of interest show a general rising trend in activation, which has been associated with global integration (Musa et al. 2025), but some pairs (notably A1 and PMC, and V1 and SPL) show higher degrees of correlation. Graph Theory analysis provides a means to formally quantify these relationships across many nodes. The insights gained from this analysis are described in the following sub-sections.

#### 4.2.1 Order-Independent Integration at the Global Level

We used global network measures and normalized them to a random network model to assess whether they differed significantly from the null model (Váša and Mišić 2022; Ghaderi et al. 2023; Musa et al. 2025). Both the combined and order-dependent networks were shown to be ‘true’, i.e., non-random networks (Fig. 7). Specifically, these networks showed significantly lower Global Efficiency, consistent with the greater constraints in information processing present in a real brain network (Latora and Marchiori 2001; Bullmore and Sporns 2009), 2), significantly higher clustering coefficients, a measure of how well the network formed clusters of highly correlated nodes, (Achard and Bullmore 2007; Tomasi et al. 2013), and significantly lower Energy reflecting the effort used to maintain organization in the network (Watts and Strogatz 1998; Rubinov and Sporns 2010).

#### 4.2.2 Network Integration: Modules and their Local Hubs

We next examined modularity, a mesoscale measure of how well a network can be divided into communities (modules), to understand how top-down auditory and bottom-up visual cues were integrated into the reach network. We also consider local ‘eigenvector centrality’ hubs within these modules, which generally aligned well with activation maps from the execution phase (Fig. 8-9).

**Module 1.** As shown in Figure 6A, the temporal-frontal-somatomotor module contained nodes in the primary auditory cortex and areas involved in reach planning execution, including premotor regions, SMA, and most of the bilateral somatomotor cortex (Hanakawa et al. 2008; Gallivan et al. 2011; Pilacinski et al. 2018). Local ‘Eigenvector’ hubs in this module included bilateral anterior cingulate cortex, implicated in the ‘action-mode network’ (Dosenbach et al. 2025). Although one cannot infer anatomy or causality from this data, based on previous anatomic and physiological studies, our results strongly suggest that the auditory instruction for the motor strategy (horizontal vs. vertical grip) was incorporated into motor planning and execution through this module. In particular, these components of this module are consistent with previous studies suggesting that the auditory-verbal-motor processing stream travels from the superior temporal cortex to the inferior frontal cortex and reaches the premotor cortex (Rauschecker 2011; Hickok and Poeppel 2015).

**Module 2**. The occipital-parietal-somatomotor module included much of the visual cortex, as well as parietofrontal areas involved in visuomotor transformation, including SPOC, IPS, superior parietal cortex, and portions of the left somatomotor cortex contralateral to the hand used. Hubs for this network were mainly found in the visual cortex, with one in the left somatomotor cortex. Again, based on the known physiology of these areas, our network data strongly suggest that the visual aspects of the task (location, shape, size, orientation of the object) were communicated through this module (Gallivan et al. 2009; Cavina-Pratesi et al. 2010; Singhal et al. 2013; Gallivan and Culham 2015). Overall, the components of this module are consistent with the classic notion of a dorsal (occipital-parietal-frontal) visual stream for visually guided action (Goodale and Milner, 1992; Goodale et al., 1994).

**Module 3**. While its modularity was not statistically significant, this community is worth noting because it contains many high-level temporal and frontal areas implicated in non-sensorimotor functions such as attention, salience, task selection, and decision-making (Rossi et al. 2009; Rens et al. 2017; Moerel et al. 2024).

**Order-Dependence**. Interestingly, more hubs appeared when we separated our data by cue order, including the frontal eye fields (FEF) within the dorsal attention network, the dorsal anterior cingulate cortex (ACC) within the ventral attention network, and nodes of both the dorsal and ventral somatomotor networks. These regions may be more strongly engaged in the later phases of integration, as frontal and motor regions are recruited for action execution. In particular, the dorsal ACC has been linked to action planning, decision making, and behavioral adaptation (Srinivasan et al. 2013; Kurkela and Ritchey 2024). However, local hubs were virtually identical in the two orders (Fig. 8B-C). This supports the notion that local hubs are relatively stable in their membership to specific modules.

The exception is that Modules 1 and 2 seemed to ‘compete’ for the left somatomotor hub, depending on which came first. This is reflected more broadly in the pattern that either Module 1 or 2 invaded the left somatomotor cortex (as if competing for right-hand dominance), depending on which cue appeared first. This seems to underlie our finding that Modules 1 and 2 were most important for decoding task order from these data.

#### 4.2.3 Lateral Integration: Betweenness Centrality

Is cue integration only achieved through feedforward processing? Another measure, Betweenness Centrality, provides the potential to identify putative hubs for lateral communication between modules. As shown in Figure 9, betweenness centrality hubs appeared in broader areas, including more sensory areas, especially when the data were separated by cue order. Important hubs were found within the control network and Somatomotor network for Module 1, within the visual central and visual peripheral networks for Module 2, and within the salience/attention and action-mode networks for the two modules (Seeley 2019; Dosenbach et al. 2025). These findings suggest that our two cue-dependent modules (one and two) communicate with each other before information converges in the somatomotor cortex. If so, integration is an ongoing process involving both lateral and feedforward converging projections.

### 4.3 Caveats and Limitations

Our findings suggest that (at the level of the BOLD signal) cue integration is distributed between communicating modules rather than concentrated in a few distinct areas, a pattern that has also been reported in previous studies (Cavina-Pratesi et al. 2010; Vossel et al. 2014; Bertolero et al. 2015; Gordon et al. 2018). However, several caveats are worth considering. First, we again acknowledge that our number of participants was low by current standards, although this is offset by the robust and reliable activation produced in our task (Fig. 3), our statistical analyses suggest that even the more modest effects (i.e. the order-dependence) were sufficiently reliable for univariate and GTA analysis, and ultimately the results made sense in terms of the previous literature.

Second, our attempts to derive specific sites for integration *within* ROIs have consistently failed, but it is always important to replicate results using other tasks and recording techniques. Third, a recent study suggested that the directionality and linearity of the BOLD response may not correlate with neural activity as well as previously thought (Epp et al. 2025). To some degree, functional connectivity is less sensitive to this problem, since it relies on the absolute value of BOLD-to-BOLD correlations. But still, all such results are correlative, so any discussion of causation, direct communication, or directionality is speculative. Fourth, the BOLD response likely represents the ‘democratic vote’ of large populations of neurons, so there is likely far more diversity and intermingling of cue signals within regions of interest revealed by our data. Finally, we only examined cortical networks; however, subcortical regions, the basal ganglia, cerebellum, and thalamus, likely also contribute to these networks (Battaglia-Mayer and Caminiti 2019).

### 4.4 Possible Clinical Implications

It is well established that damage to the dorsal visual stream (corresponding to our Module 2) interrupts the visually guided movement (Goodale and Milner 1992; Pisella et al. 2000; Khan et al. 2005; Khan et al. 2007) and based on our data, one would expect damage to our Modules to degrade the ability to use verbal commands for action (Wheaton and Hallett 2007). This does not mean that these modules are fixed; indeed, we expect that specific cues are flexibly integrated via distinct modular streams, depending on both the sensory modality and the cue’s relevance to the task. For example, one might speculate that switching the meaning of our cues (i.e., visual for motor strategy and auditory for location) or pairing a completely different sensory modality (e.g., somatosensory) with a different outcome (e.g., to make an eye movement) would result in an entirely different network (Groh and Werner-reiss 2002; Rens et al. 2017).

This combination of network distribution and modular specificity could explain why some patients retain partial function after focal damage, as alternative routes are used to maintain communication across the network. And again, these alternative networks might include subcortical pathways (Grefkes and Fink 2011; Battaglia-Mayer and Caminiti 2019; Guggisberg et al. 2019). Extending our task-related functional network approach could lead to a better understanding of how plasticity contributes to brain injury recovery and how degeneration in brain diseases like dementia and Parkinson’s disease can be delayed (Bedford et al. 2025; Hao and Zou 2025; Jahan et al. 2026).

### 4.5 Conclusion

Overall, these analyses suggest that multimodal integration of visuospatial cues and verbal instructions for reach is a network phenomenon, involving two separate (occipital-parietal and temporal-frontal) modules that communicate via lateral connections and converge in the somatomotor cortex. Within these modules, local network hubs may participate in this process both through relatively fixed sensorimotor roles for a given task, whereas more distributed sensory and motor ‘betweenness’ hubs may underlie the lateral communication. Finally, in our task, varying cue order had relatively little effect on cortical networks at the global level, but produced subtle, decodable effects at the mesoscale (modular) and local levels. This approach, combining event-related recordings, univariate analysis, and whole-brain functional connectivity, may be also useful for understanding cortical networks for rehabilitation following damage. Overall, this study shows that multisensory-motor integration of top-down instructions and bottom-up sensory cues is achieved through parallel but laterally communicating and serially converging modules.

## Article Information

### Data and Code Availability statement

Data and code are available for requests that comply with federal data sharing policies, pending approval from the requesting researcher’s local ethics committee and agreement on how to share author credits on any resulting publication.

### CRediT statement

GNL: Data curation, Formal Analysis, Conceptualization, Visualization, Writing - Original draft, Writing - review and editing. AL: Methodology, Investigation, Writing - review and editing. AG: Formal Analysis, Conceptualization, Writing - review and editing. LM: Formal Analysis, Writing - review and editing, SM: Conceptualization, methodology, Writing - review and editing. EF: Conceptualization, Methodology, Writing - review and editing. JDC: Conceptualization, Methodology, Writing - review and editing, Funding acquisition, Project administration, Supervision. **Funding statement**: This work was supported by grants from the Canadian Institutes of Health Research (grant number RGPIN-2022-04527) and the Natural Sciences and Engineering Research Council of Canada (grant number MOP-68812). Luabeya and Ghaderi were supported by the Vision: Science to Applications Program, funded in part by the Canada First Research Excellence (CFREF) program. Le was supported by the National Science and Engineering Research Council Canada Brain in Action CREATE Program. Crawford was supported by a Canada Research Chair.

### Ethics approval statement

This experiment was approved by the York University Human Participants Review Committee (Certificate #: 2015-219; initially approved July 30, 2015, and renewed November 22, 2021).

## Acknowledgements

The authors thank Dr. Xiaogang and Saihong Sun for technical support and Dr. Jessica Parker, Dr. Cristina Rubino and Petros Georgiadis for comments and proofreading the manuscript.

