## Supplementary Figure 1 for "Modular Integration of Auditory Instructions and Visual Cues into the Cortical Reach Network"

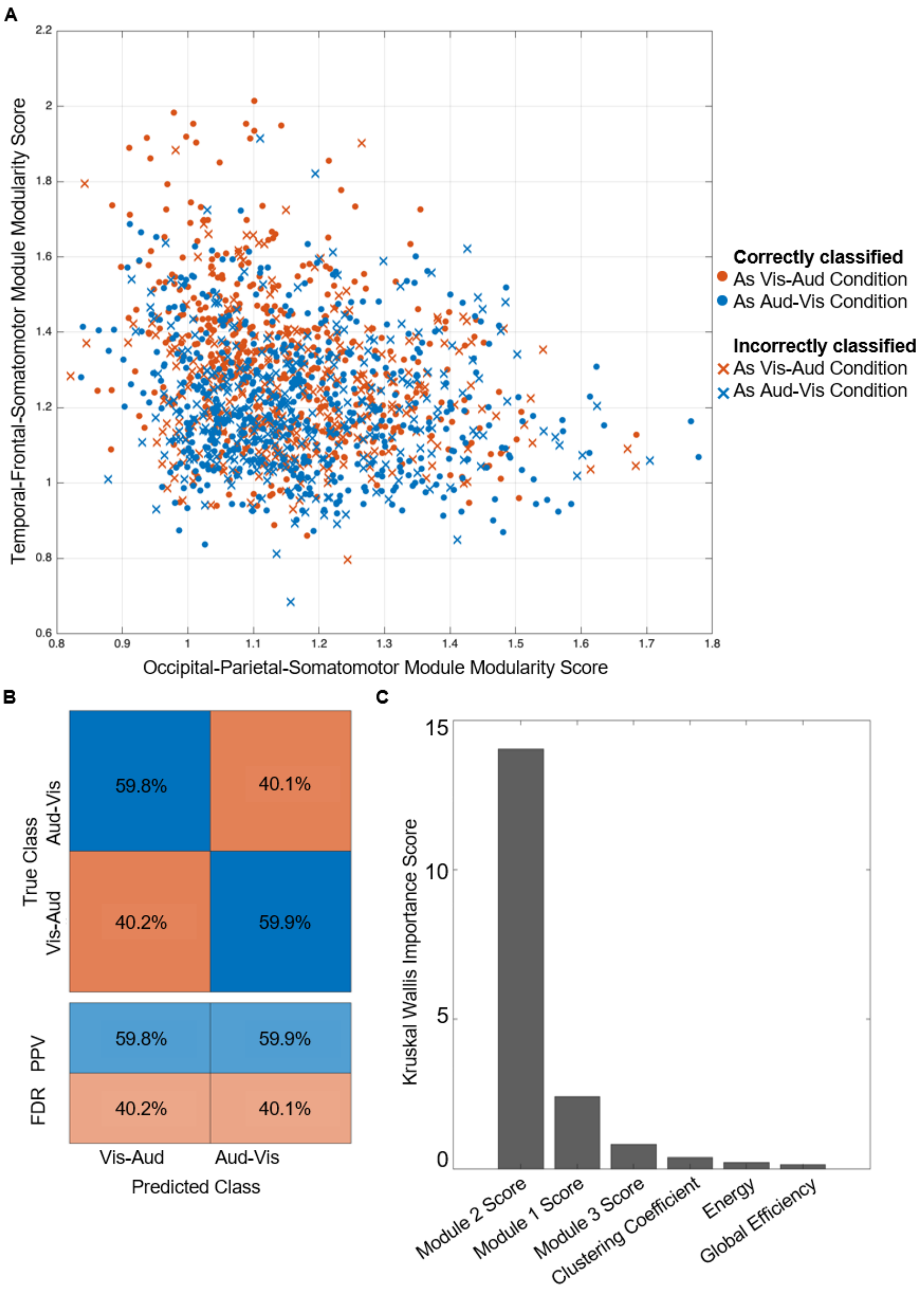


Supplementary Figure 1. Classifier Analysis represents the performance of the Narrow Neural Network classifier to differentiate between the Visual-Auditory Order (Vis-Aud) and Auditory-Visual Order (Aud-Vis). A) Scatterplot showing the distribution of modularity values from the top two predictors based on the Kruskal Importance Score (Module 2: Occipital-Parietal-Somatomotor Module and Module 1: Temporal-Frontal-Somatomotor Modularity Index). A circle represents a correctly predicted trial order, and a cross represents an incorrectly predicted trial order. B) displays the confusion matrix, comparing the predicted data point identity (Vis-Aud or Aud-Vis) with its true identity. The bottom panel table shows the positive predictive values (PPV) and false discovery rates (FDR)

of the Grasp data (left column) and the Place data (right column). C) The Kruskal-Wallis importance scores of each feature were used to classify the order of cue presentation. A higher value indicates greater separation between classes based on that feature – meaning the feature is more useful for classification.
